# Cytoplasmic localization of 5hmC reveals RNA hydroxymethylation in breast cancer

**DOI:** 10.64898/2026.10.21.760667

**Authors:** Jinrui Wen, Lisa Del Bel Belluz, Silvia Ottaviani, Fotini Poyia, Hemeng Yu, Saachi Sachdev, Liliane El Eid, Katherine Garner, Krina Mistry, Emily Hayes, Nina Moderau, Pramodh Chandrasinghe, Justin Stebbing, Biancastella Cereser

**Author notes:** These authors contributed equally to this work as co-senior authors (Justin Stebbing, Biancastella Cereser). Corresponding author: (BC).

## Abstract

DNA methylation (5-methylcytosine, 5mC) is a major repressive epigenetic mark that can be iteratively oxidized by ten-eleven-translocation (TET) enzymes to 5-hydroxymethylcytosine (5hmC); beyond DNA, modifications of RNA can also affect transcript stability and processing, yet RNA hydroxymethylation remains unexplored in breast cancer (BC). Here, we identify a quantifiable cytoplasmic 5hmC signal in BC cell lines and, more prominently, in human BC tissue, using immunoassays and digital pathology, and confirm with an adapted, RNase-sensitive, RNA-specific ELISA that this signal reflects hydroxymethylation of total RNA and mRNA. Cytoplasmic 5hmC was significantly enriched in triple-negative (ER-/PR-/HER2-) and grade 3 tumors, and correlated with increased cytoplasmic TET2/TET3 in tissue and *TET3* upregulation in TCGA. As a leading candidate mechanism, *TET3* expression was associated with RNA-processing/spliceosomal transcripts, and *TET3* knockdown in three TNBC cell lines reduced a subset of these targets, independently supported across 70 DepMap breast cancer lines.

## Background

Breast cancer (BC) is a heterogeneous disease, with distinct molecular subtypes that show different clinical behaviors and treatment responses. Several studies have demonstrated how epigenetic alterations, in particular DNA methylation, play a critical role in BC development and progression and may also be used as biomarkers to classify the BC tumor subtypes ^1–4^. While there has been a focus on modifications involving 5-methylcytosine (5mC), in the past decade the discovery of 5-hydroxymethylcytosine (5hmC) and the related activity of TET enzymes (TET1, TET2, and TET3) has facilitated the development of further biomarkers and therapeutic targets for BC. Global 5hmC levels are significantly decreased in many human cancers, including those without mutations in *TET* or *IDH* genes, the latter being essential for the production of 2-oxoglutarate (2OG) and required for the activity of the TET enzymes ^5^. The global loss of 5hmC ranges from ∼50% to over 90% in tumors of almost all major histological types, including lung ^6^, colorectal ^7^, pancreatic ^8^, prostate ^9^, gastric ^10^, liver ^6^, melanoma ^11,12^, glioma ^13^, and genitourinary tumors ^14^.

Previous studies using dot blots and commercial immunoassays have successfully identified 5hmC in DNA from various cell lines and tissues, including breast tissue, at varying concentrations^15,16^. Additionally, these assays effectively detect 5hmC in the DNA of colon and breast cell lines, where the estimated content is reported to be less than <0.02% of total DNA ^15^. However, these assays do not offer insights into the localization of 5hmC outside the cell nucleus, where different classes of RNA and mitochondrial DNA could present potential targets for 5hmC.

In several cancer types, modifications by 5mC RNA methyltransferases, observed in mRNA, tRNA, rRNA, lncRNA, and miRNA, can affect stability, processing, cleavage, transcription, translation, and ultimately the regulation of various cellular functions ^17^. RNA 5mC modification has gathered attention in solid tumor research, with the writer NSUN2 (a methyltransferase catalyzing the methylation of cytosine to 5mC in various RNAs) implicated in pathways promoting cell progression, migration, and invasion in BC and other cancers ^18^. Likewise, RNA hydroxymethylation can also regulate mRNA translation and stability during embryonic stem cell (ESC) differentiation ^19^ and in mouse models of chronic environmental stress ^20^, while in glioma, NSUN5 induces recruitment of TET2 from the cytoplasm to chromatin to induce hydroxymethylation and ultimately degradation of *CTNNB1* mRNA ^21^.

In mouse liver tissue, 5hmC mainly occurs in mRNA rather than in other classes of RNA, and this modification is significantly lower in human colorectal cancer and hepatocellular carcinoma samples compared to adjacent controls ^22^. However, while 5hmC has also been previously detected using LC-MS/MS/MS in RNA samples from mouse brain, heart, pancreas, and spleen, and in vitro in cervical cancer cells (HeLa) and melanoma (WM-266–4) cells ^23^, 5hmC has not been previously reported in the RNA of breast tissue or cell lines.

In this study, we aimed to investigate the cytoplasmic localization of 5hmC in breast cancer cells, demonstrate the presence of RNA hydroxymethylation, and analyze the correlation of 5hmC levels with key tumor characteristics.

## Results

### Cytoplasmic localization of 5hmC in BC cells reflects hydroxymethylation of the RNA

To characterize the distribution of 5hmC within breast cancer cells, we performed immunofluorescence staining across four commonly used BC cell lines (BT474, MCF7, MDA-MB-231, and SKBR3). In all cases, 5hmC was clearly detectable in both nucleus and cytoplasm (**Fig. 1A**). Across all four cell lines, mean 5hmC fluorescence intensity was significantly higher in the nucleus than in the cytoplasm (linear mixed-effects model, LMM: log₂(intensity) ∼ Class + (1|Replicate); BT474: estimate = 0.678, p = 4.87 × 10⁻¹⁶; MCF7: estimate = 0.673, p = 2.75 × 10⁻⁶; MDA-MB-231: estimate = 0.325, p = 2.15 × 10⁻⁵; SKBR3: estimate = 0.360, p = 2.61 × 10⁻¹⁰; **Fig. 1B** and **Supplementary Table 2, Supplementary Data 1**), with median nuclear intensities ranging from 46.3 (MDA-MB-231) to 163 (MCF7). However, when total 5hmC signal was considered by accounting for compartment size via integrated density, the cytoplasmic fraction of total cellular 5hmC signal ranged from 44.3% to 51.7% across cell lines, with none significantly different from equal distribution (LMM tested vs 0.5; all n.s., **Fig. 1C**). This indicates that, despite its lower 5hmC concentration, the cytoplasm’s larger volume compensates such that it harbours a roughly equal proportion of total cellular 5hmC signal, a finding consistent with a biologically meaningful cytoplasmic pool of hydroxymethylated nucleic acids. Nuclear/cytoplasmic (N/C) intensity ratios confirmed nuclear enrichment of 5hmC concentration across all lines (**Fig. 1D**), reaching statistical significance in MCF7 (LMM estimate = 0.652, p = 0.038) and SKBR3 (LMM estimate = 0.348, p = 0.020). Notably, MDA-MB-231 cells, which represent a triple-negative/basal-B subtype, displayed the lowest N/C ratio among the four lines (median = 0.141) and were the only cell line in which nuclear enrichment of 5hmC concentration did not reach statistical significance (LMM estimate = 0.341, p = 0.330), suggesting a more equal nuclear–cytoplasmic distribution in this subtype. This observation is consistent with the more prominent cytoplasmic 5hmC enrichment observed in triple-negative breast cancer tissues described below, though between-cell-line differences in N/C ratio did not reach statistical significance overall (Kruskal-Wallis, p = 0.223).

**Fig. 1.**
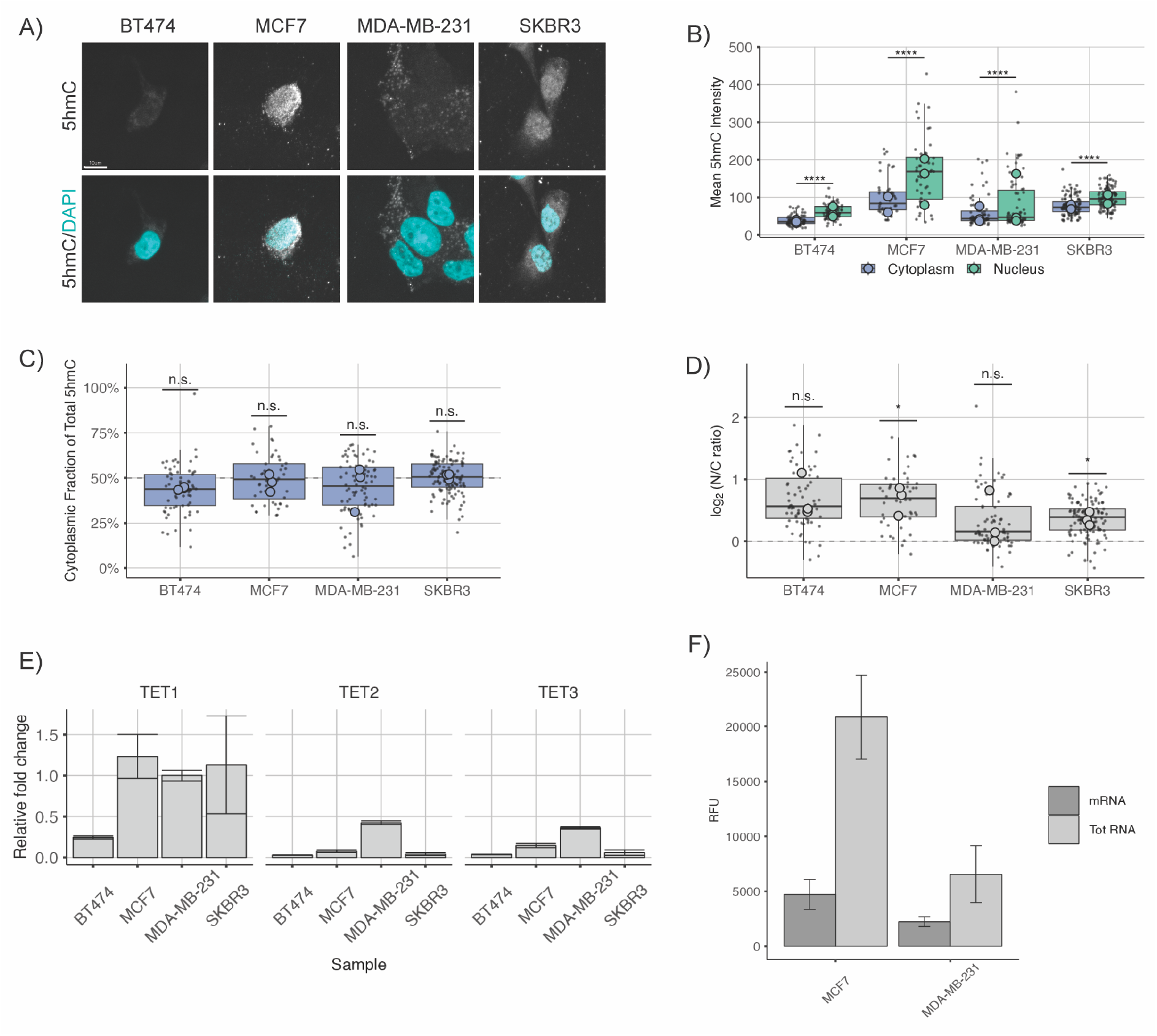
Quantitative analysis of 5hmC content and TET expression in four BC cell lines. (A) Representative immunofluorescence images show 5hmC signal (greyscale) and DAPI nuclear staining (cyan) across four BC cell lines. (B) 5hmC mean fluorescence intensity is higher in the nucleus than in the cytoplasm across all cell lines (linear mixed-effects model, log2(intensity) ∼ Class + (1|Replicate), per cell line). (C) When accounting for compartment size via integrated density, the cytoplasmic fraction of total 5hmC signal ranged from 44.3% to 51.7% across cell lines, with none significantly different from equal distribution (LMM tested vs 0.5; all n.s.), indicating that the cytoplasm’s larger volume compensates for its lower 5hmC concentration. (D) Log2 N/C intensity ratios confirm nuclear enrichment of 5hmC concentration, reaching significance in MCF7 and SKBR3 (LMM tested vs 0; per cell line). Boxplots in B–D display the IQR and median; jittered dots show individual cells; large outlined points represent biological replicate medians (n = 3). (E) RT-qPCR shows variable *TET1*, *TET2* and *TET3* expression across cell lines, shown as fold changes relative to *TET1* in MDA-MB-231 (SD, n = 3). (F) 5hmC is detectable in both total RNA and mRNA fractions in MCF7 and MDA-MB-231 cells (MethylFlash™ adaptation, RFU, SD, n = 3). Panels E–F are descriptive and do not include significance testing. Significance: * p < 0.05, ** p < 0.01, *** p < 0.001, **** p < 0.0001; n.s., non-significant.

These findings indicate that 5hmC amount and distribution do not directly reflect the pattern of expression of the three TET enzymes in the same cell lines (**Fig. 1E** and **Supplementary Table 2, Supplementary Data 1**), consistent with previous publications (16).

The identification of cytoplasmic 5hmC within cancerous tissue implies an enrichment of 5hmC extending beyond nucleic acids confined to the cell nucleus. To explore potential targets responsible for the cytoplasmic enrichment of hydroxymethylation, we employed commercial kits, originally established for the detection of 5mC in the RNA and adapted for the detection of 5hmC, on total RNA and on mRNA extracted from two BC cell lines (MCF7 and MDA-MB-231). Hydroxymethylation is a feature of both total RNA (short and long RNA, including mRNA) and mRNA specifically in both cell lines.

The derived relative signal showed a significant difference (p = 0.0004) of 5hmC in the total RNA between MCF7 and MDA-MB-231 cells (**Fig. 1F**), but no significant difference in the levels of 5hmC in the mRNA only (p = 0.617). While a positive control comprising 5hmC in DNA is a necessary technical control for the adapted kit (**Supplementary Fig. 1, Supplementary Data 1**), the different binding ability of DNA to an assay designed for RNA makes it unsuitable for the precise quantification of 5hmC across different RNA classes. This highlights the necessity for the development of controls involving RNA with a predefined percentage of 5hmC.

To confirm that this signal originated from RNA rather than contaminating DNA, we performed a series of validation experiments on the adapted ELISA (**Fig. 2**). Bioanalyzer electropherograms of the RNA samples used confirmed high RNA integrity (RIN ≥ 9 in all samples) with distinct 18S and 28S rRNA peaks and no detectable genomic DNA contamination (**Fig. 2A**). An RNA dilution series using DNase-treated total RNA (300, 150, 75 and 0 ng input) showed a dose-dependent, linear decrease in signal (R² = 0.93, p = 0.038), consistent with a quantitatively reliable, RNA-derived signal rather than non-specific background or DNA contamination (**Fig. 2B**). Treatment of RNA samples with RNase I abolished the 5hmC signal, reducing it to levels comparable to the no-RNA control, confirming that the signal is RNA-dependent (**Fig. 2B**). Finally, serial dilutions of 5hmC-DNA positive controls exhibited poor, non-linear binding with high inter-replicate variability on the RNA-capture plate, demonstrating that the assay is not optimized for DNA detection and confirming its RNA-specific design (**Fig. 2C**). Together, these controls indicate that the cytoplasmic signal described above, and the signal detected in extracted total RNA and mRNA, reflect genuine RNA hydroxymethylation rather than contaminating DNA.

**Figure 2.**
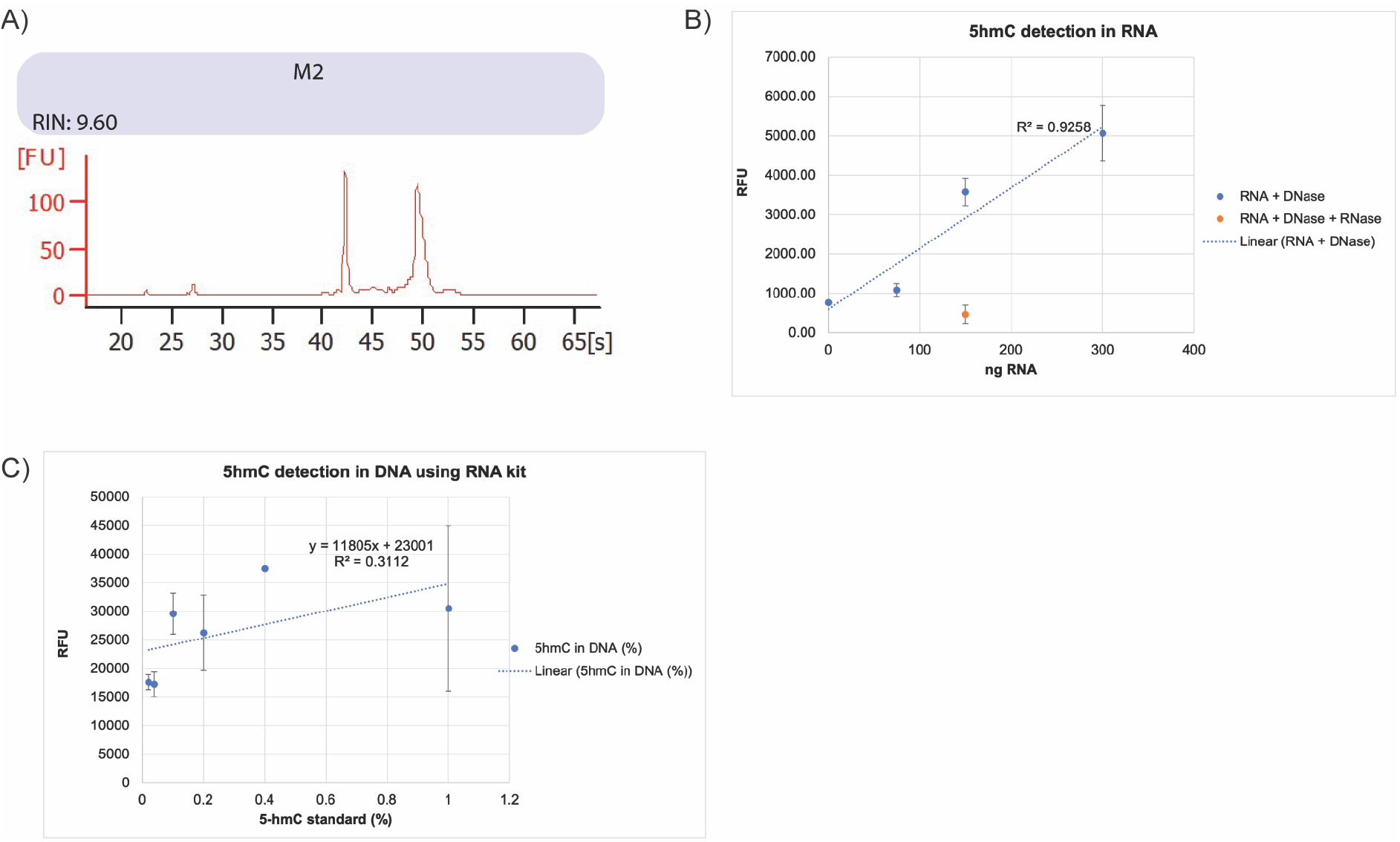
Validation of the adapted 5hmC RNA ELISA assay. (A) Bioanalyzer electropherogram of DNase-treated total RNA showing high integrity (RIN ≥ 8.5) and absence of genomic DNA peaks. (B) RNA dilution series (300–0 ng; blue) shows linear signal decrease (R² = 0.95, p = 0.038). RNase I-treated RNA (orange) yields background signal, confirming RNA dependence. (C) 5hmC-DNA standards show non-linear, variable binding on the RNA-capture plate, confirming RNA specificity and DNA exclusion.

### Sublocalization of 5hmC to the cytoplasm of BC tissue

Global levels of 5hmC are reduced in human cancers, including in BC tissues, compared to the matching healthy counterparts ^24^; however, the distribution of 5hmC levels within a tumor and cancer cells in patients is still unknown. This prompted us to investigate the cellular localization of 5hmC within BC cells in human tissue samples. Immunohistochemistry (IHC) was performed on the cohort of 43 samples from matching IDC and adjacent phenotypically normal tissue (**Fig. 3A** and **Supplementary Table 5, Supplementary Data 1**). Compared to the adjacent tissue, 5hmC levels in the nuclei of BC cells were not significantly different (Welch Two Sample t-test, p=0.535, **Fig. 3B**). However, levels of 5hmC were significantly higher in the cytoplasm of cancer cells, compared to the normal counterpart (Welch Two Sample t-test, p=0.003, **Fig. 3B**). An average of approximately 50% of cancer cells show an enrichment of 5hmC in the cytoplasm, compared to an average of approximately 21% of normal breast cells. A paired analysis is particularly useful when comparing these samples, as it helps to account for individual differences. A matched analysis confirmed the higher percentage of positive cytoplasmic levels of 5hmC in the cancer cells compared to the non-affected cells (n = 20 sample pairs, paired t-test, p=0.003, **Fig. 3B**).

**Fig. 3.**
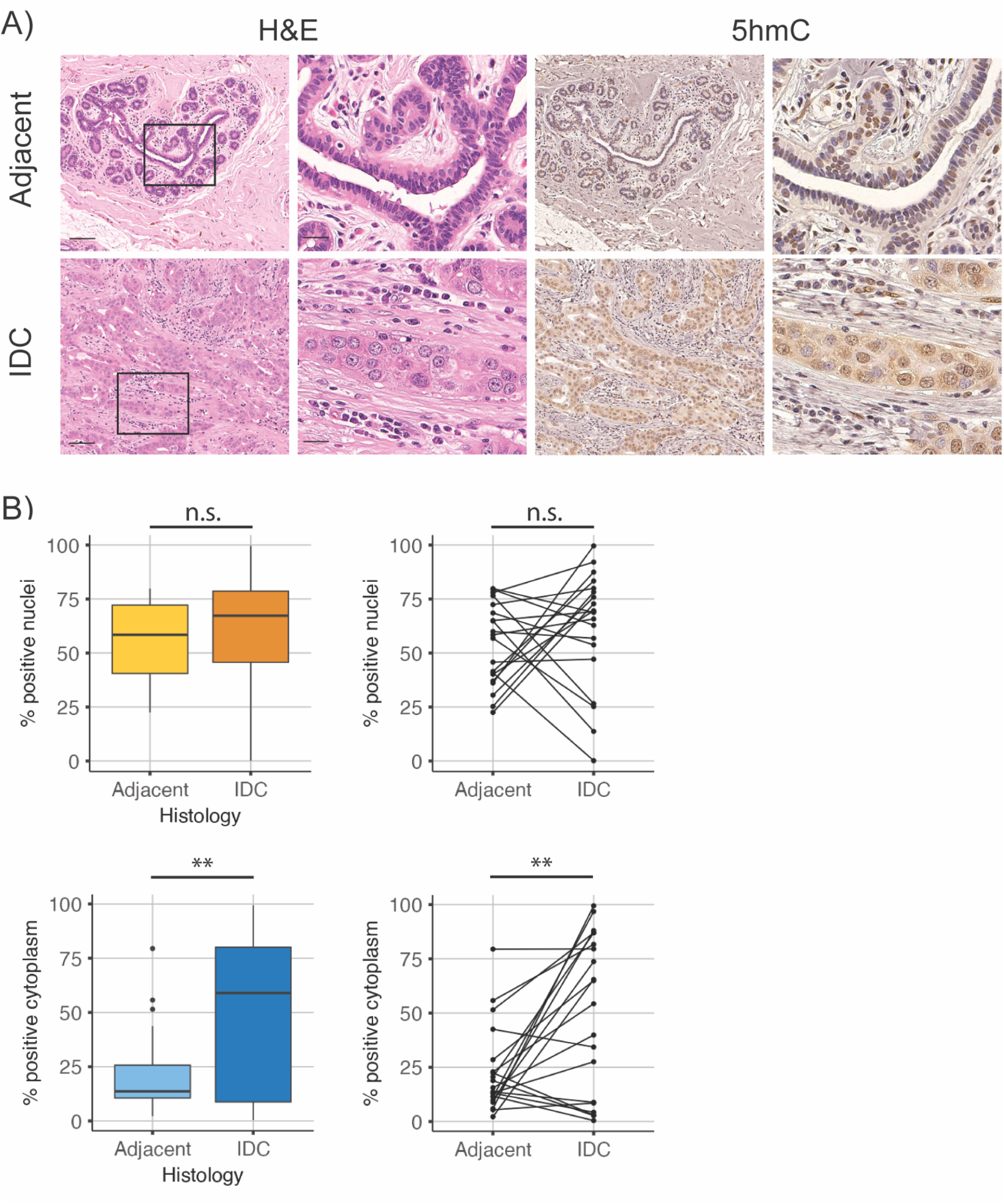
Levels of 5hmC are higher in the cytoplasm of BC tissue. (A) H&E 5hmC immunohistochemistry of representative serial sections of normal adjacent breast tissue and the matching invasive ductal carcinoma. High-power images are shown in X) – Y). Scale bar=100μm; inset scale bar=20μm. (B) Unpaired and paired analysis shows that the cytoplasm of cancer cells contains significantly higher levels of 5hmC compared to the normal adjacent counterpart. For unpaired graphs, n=23 for adjacent and n=24 for IDC. For paired graphs, n=20 pairs. Significance is indicated as ** (p < 0.01) or n.s. (non-significant). Boxplots display the interquartile range (IQR) and median.

As an experimental control to assess the specificity of cytoplasmic staining, we evaluated a subset of 12 breast samples (6 cancer sections and 6 matched non-affected adjacent tissues) for abundance of 5mC (**Supplementary Fig. 2A, Supplementary Data 1**). Despite variations among the samples, nuclear levels of 5mC in cancer samples were not significantly elevated compared to adjacent samples (Welch Two Sample t-test, p=0.477, **Supplementary Fig. 2B, Supplementary Data 1**), and both tissue types exhibited high levels of positivity (>70% positive cells on average). In contrast, less than 3% of cells in the cytoplasm were detected as positive for 5mC levels. Although the difference in 5mC content between cancer and non-affected cells may appear statistically significant (p = 0.004), these results should be interpreted with caution due to the low percentage, which may be subject to quantification errors.

It is known that global 5hmC levels in DNA vary markedly among normal human tissues, with brain tissue exhibiting significantly higher levels of 5hmC compared to breast, liver, testes, placenta, colon, or blood DNA ^15,16,25^. To investigate whether the cytoplasmic sublocalization of 5hmC in cancer cells is a unique characteristic of mammary epithelium cells as opposed to other tissues, for a further control 5hmC levels were assessed in a cohort of 5 patients affected by colitis-associated cancer tissues (8 samples), along with matched non-malignant epithelium (quiescent colon, 12 samples). Interestingly, this assessment confirmed significantly higher levels of cytoplasmic 5hmC exclusively in colonic cancer cells compared to the quiescent colon (Welch Two Sample t-test, p=0.016) (**Supplementary Fig. 3, Supplementary Data 1**). These findings indicate that the cytoplasmic distribution of 5hmC is not restricted to BC, suggesting diverse mechanisms of epigenetic regulation based on hydroxymethylation in other healthy and cancerous tissues. Due to each quiescent sample being primarily composed of transverse crypt sections, visualizing a sufficient number of nuclei in apical colonic mucosal cells for 5hmC enrichment assessment, as observed in previous studies ^25^, was not feasible.

### Levels of cytoplasmic 5hmC correlate with differences in the hormonal receptor status of the cancer and grade of the tumor

Previous studies have further shown a correlation between global 5hmC levels and the hormonal receptor status of BC specimens and highlighted an association between low 5hmC levels and a lower rate of survival ^26,27^. To investigate whether a similar association could be detected based on levels of cytoplasmic 5hmC, the sample cohort was stratified based on the routinely used clinical subsets according to the expression of estrogen receptor (ER), progesterone receptor (PR), and human epidermal growth factor receptor 2 (HER2) in the cancer specimen (**Fig. 4**). Matched analyses indicated that cytoplasmic levels of 5hmC were significantly elevated only in the cancer cells of triple-negative (ER-/PR-/HER2-) patients (Welch Two Sample t-test, p=0.012, and, after matched analysis on n = 9 sample pairs, paired t-test, p=0.008, **Fig. 4** and **Supplementary Table 5, Supplementary Data 1**). Similarly, we stratified the samples based on the grade of the tumor and identified a significant increase in the levels of cytoplasmic 5hmC in the cancer cells of patients diagnosed with Grade 3 BC, independently of the subtype, compared to their adjacent controls in the entire cohort (Welch Two Sample t-test, p=0.010) and after matched analysis (paired t-test, p=0.008) (**Fig. 4** and **Supplementary Table 5, Supplementary Data 1**).

**Fig. 4.**
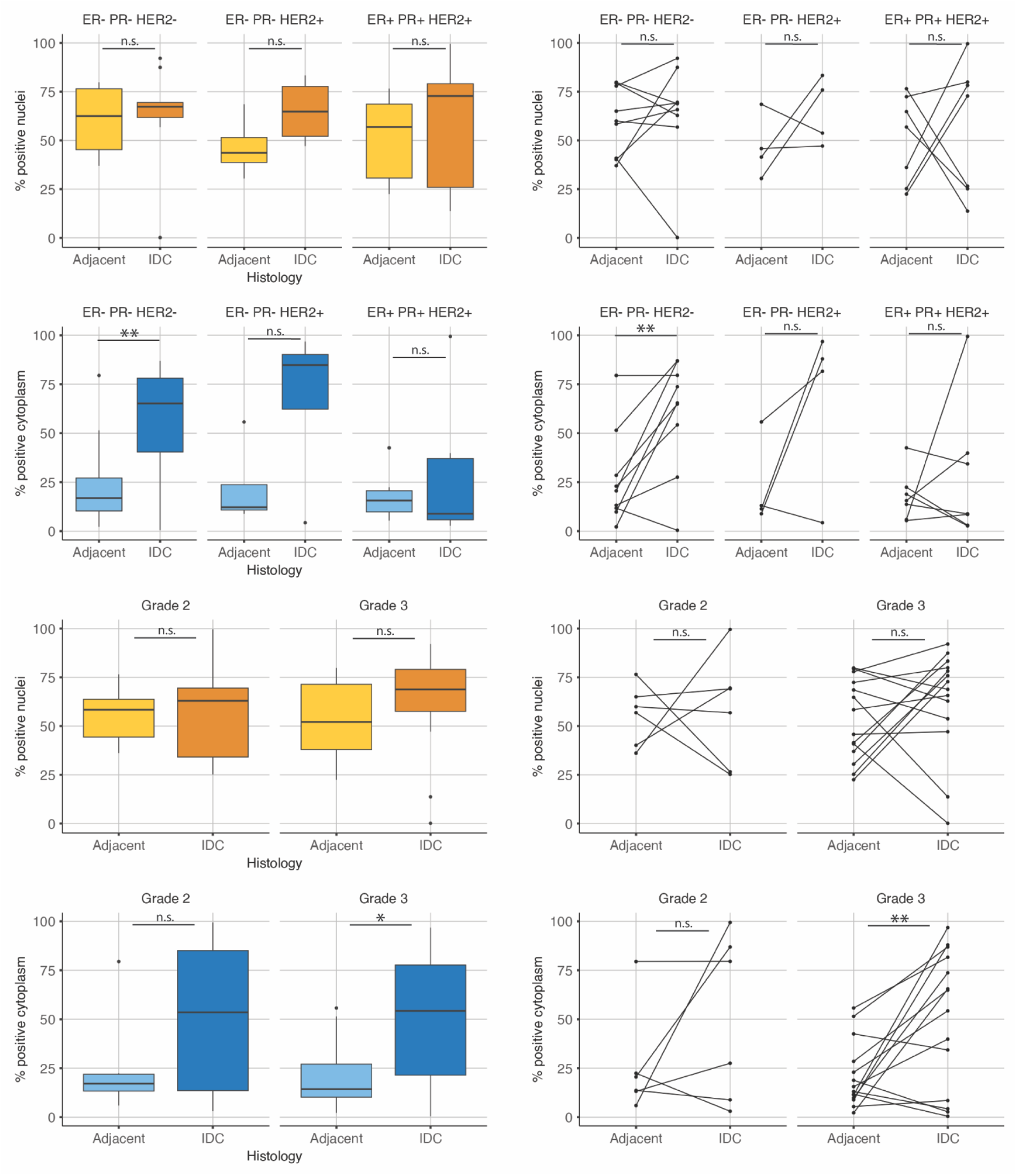
Cytoplasmic levels of 5hmC are elevated in triple-negative BC samples. The graphs show the abundance of 5hmC (as a percentage of positive cells) in the nuclei or the cytoplasm of BC samples, stratified by the receptor status or the grade of the cancer section. Ordinary one-way ANOVA and paired t-tests, respectively, were performed only on the samples where cancer and matched normal (non-affected) epithelium were provided. For sample size, see **Supplementary Information**. Significance is indicated as * (p<0.05), ** (p < 0.01) or n.s. (non-significant). Boxplots display the interquartile range (IQR) and median.

### Cytoplasmic TET2 and TET3 expression is significantly higher in BC epithelium of ER-/PR-samples than in the non-affected tissue

The cytoplasmic 5hmC enrichment described above raises the question of which enzymatic machinery could explain it; as the only known catalysts of 5hmC formation, we therefore investigated TET1, TET2 and TET3 as candidate drivers of this phenomenon in BC tissue. All three members of the TET family (TETs) can catalyze the conversion from 5mC to 5hmC. However, our *in vitro* results suggest that there may not be a clear link between the expression of a specific *TET* mRNA and the 5hmC content in the cytoplasm of BC cells. Previous studies have however emphasized the prominent role of TET2 in BC ^28,29^. To investigate the association between TET proteins and the cytoplasmic localization of 5hmC in BC tissues, particularly in ER-/PR-samples, we first assessed the expression and localization of TET1, TET2, and TET3 via immunohistochemistry in a cohort of 11 ER-/PR-breast samples (**Fig. 5A-B**). Different expression patterns of the three proteins were observed in the breast epithelium. While the nuclear expression of all TETs was comparable in both non-affected and malignant breast tissues (one-way ANOVA, p=0.566), TET2 and TET3 expression was significantly higher in the cytoplasm of cancer cells compared to non-affected controls (unpaired t-test, for TET2 p=0.018; for TET3 p=0.027, **Fig. 5B** and **Supplementary Table 5, Supplementary Data 1**). This mirrors the significant difference in cytoplasmic 5hmC levels mentioned earlier in the same samples. In the subset of ER-/PR-samples with available paired control-cancer data (n=5 pairs), a matched analysis further demonstrated a significant elevation in the cytoplasmic expression of both TET2 and TET3 (paired t-test, for TET2 p=0.023; for TET3 p=0.027, **Fig. 5B** and **Supplementary Table 5, Supplementary Data 1**), in accordance with the significant difference in 5hmC levels between IDC and adjacent matched tissue in this patient group.

**Fig. 5.**
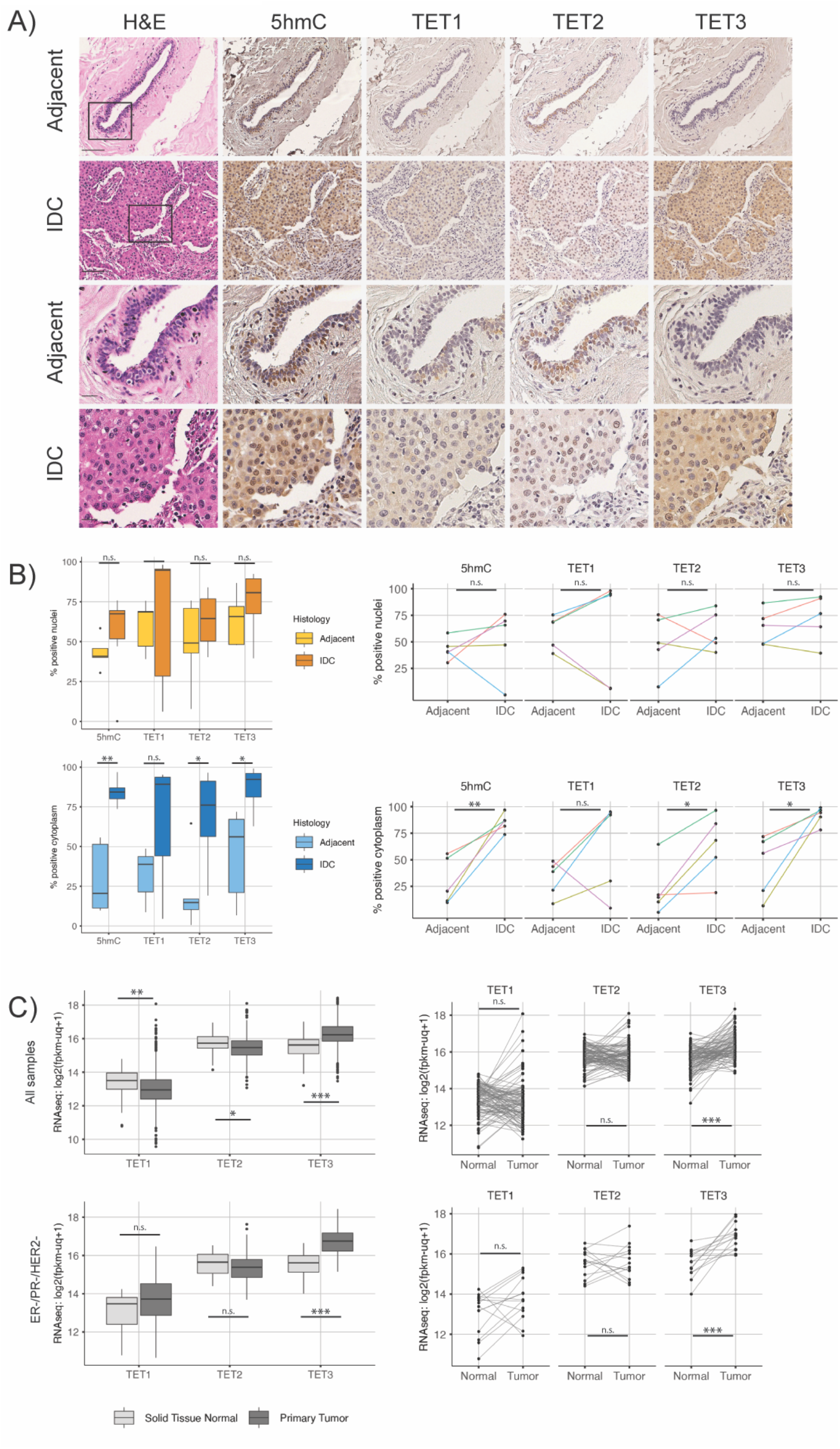
Elevated TET2 and TET3 expression in the cytoplasm of ER-/PR-BC samples. (A) H&E and immunohistochemistry staining for 5hmC, TET1, TET2, and TET3 in serial sections of normal breast tissue and corresponding invasive ductal carcinoma. On the bottom two rows, high-power images from zoomed areas highlight protein localization. Scale bar=100μm; inset scale bar=25μm. (B) Paired and unpaired analysis of the quantification of IHC demonstrates significantly elevated TET2 and TET3 levels in the cytoplasm of cancer cells compared to normal tissue. (C) A significant increase in *TET3* mRNA expression is found in the cancer samples compared to the non-affected tissues, in both unpaired and paired analyses from the TCGA dataset, in the total sample cohort and in the restricted subset of triple-negative BC. Boxplots display the interquartile range (IQR) and median. Significance is indicated as * (p<0.05), ** (p < 0.01), ***(p < 0.001) or n.s. (non-significant).

### *TET2* and *TET3* overexpression is associated with altered RNA processing and protein–RNA complex pathways in BC

To validate the dataset across a larger sample size, UCSC Xena was used to mine TCGA data from the GDC BRCA cohort, and to extract mRNA expression levels of the TET genes in a subset of samples of invasive ductal carcinomas (n = 823 primary tumors and 102 normal tissue). In these unpaired comparisons, TET3 showed the strongest increase in tumor tissue (p < 0.0001, **Fig. 5C**, **Supplementary Fig. 4A** and **Supplementary Table 5, Supplementary Data 1**), whereas TET1 and TET2 did not exhibit consistent differences between tumors and normals. While there was no significant difference between the expression levels of TET2 and TET3 in normal tissues (p = 0.858), paired analysis revealed a significantly higher expression of exclusively TET3 in BC samples compared to normal adjacent tissues (p < 0.0001, **Fig. 5C** and **Supplementary Fig. 4B, Supplementary Data 1**). Further stratifying the TCGA samples based on their receptor status and conducting paired samples analysis revealed notable differences in the expression levels of TET mRNAs across subtypes. Specifically, TET3 expression was significantly elevated in triple-negative cases (ER-/PR-/HER2-, unpaired t-test p < 0.0001 and paired t-test p < 0.001, **Fig. 5C** and **Supplementary Table 5, Supplementary Data 1**), consistent with the increased protein expression of TET3 observed in the smaller subset of samples using IHC, as described above. Interestingly, TET3 expression was significantly elevated across the whole TCGA cohort (**Supplementary Fig. 4** and **Supplementary Table 5, Supplementary Data 1**).

To understand the potential implications of differences in *TET3* expression on gene expression in breast cancer (BC), we performed a differential expression analysis on a subset of matched cancer-normal samples, determining differences in expression among samples expressing each TET mRNA. A cohort of breast ductal carcinoma and matched normal tissue samples (n = 101 patients) for which gene expression data was available (BCD cohort) was stratified based on the normalized expression levels of the three TET genes in cancer tissues relative to their matched normal tissues. "TET-high" cancer samples indicate those in which TET gene expression was higher in cancer tissues compared to the matched normal tissues, whereas "TET-low" samples indicate higher TET gene expression in normal tissues than in cancer tissues.

DESeq analysis revealed significant variations in gene expression associated with each TET gene. In the TET1-high group, out of 26,011 expressed genes, 24% (n = 6158 genes) were significantly upregulated (p adj < 0.5) and 4.9% (n = 1270) were significantly downregulated. In the TET2-high group, 26% (n = 6633) showed significant upregulation, while 7.4% (n = 1924) showed significant downregulation. Finally, in the TET3-high group, 23% (n = 5837) were significantly upregulated, and 7.7% (n = 2002) were significantly downregulated (**Fig. 6A-B** and **Supplementary Fig. 5A, Supplementary Data 1**).

**Fig. 6.**
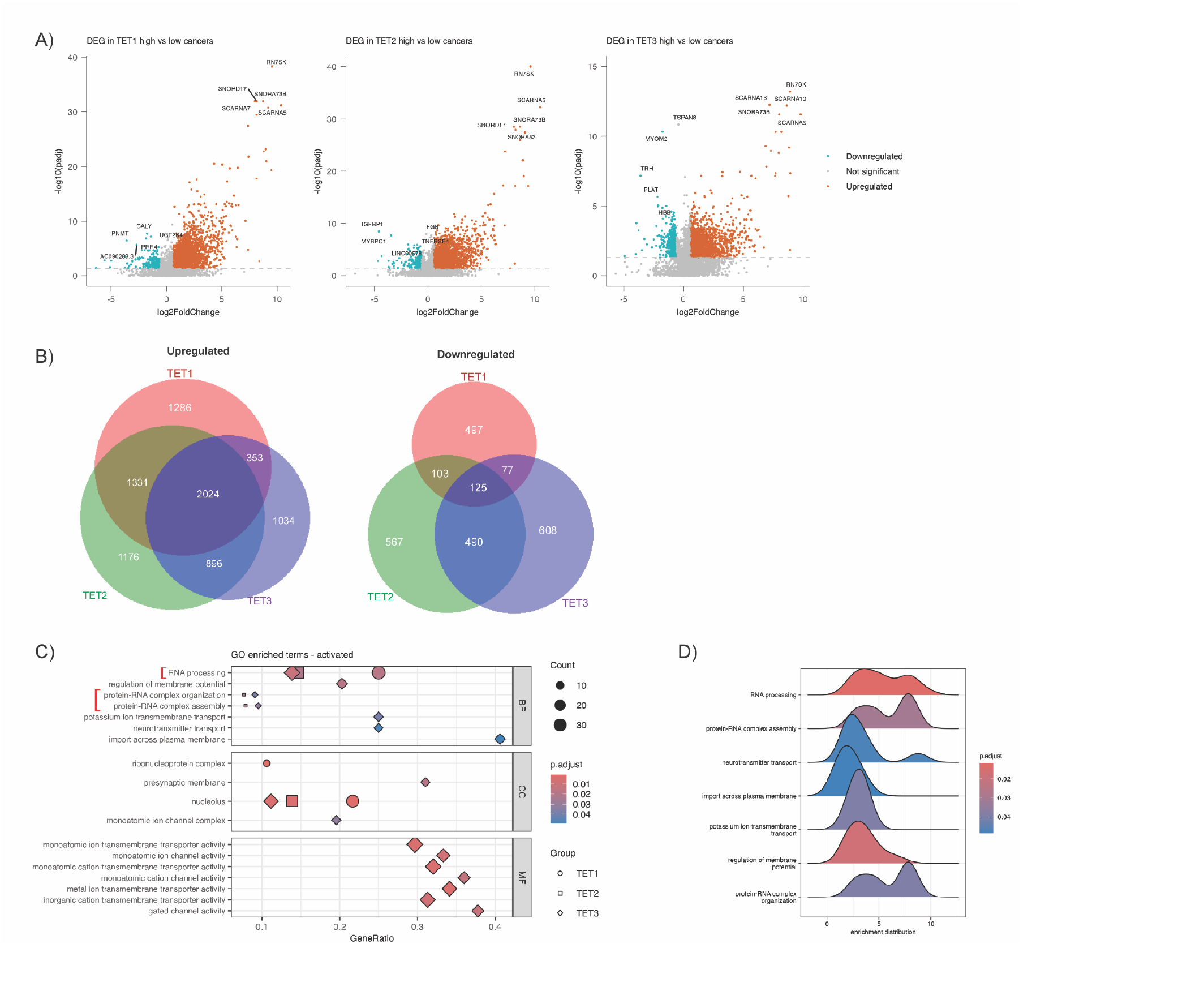
Gene Set Enrichment Analysis associated with each TET in breast ductal carcinoma. (A)Volcano plots showing DEG results for each TET group. The x-axis indicates log2 fold change in gene expression, while the y-axis represents the -log10 of the adjusted p-values. Significant upregulated DEGs (p adj < 0.05, |logFC| > 0.6) are shown in orange, and significant downregulated DEGs (p adj < 0.05, |logFC| > 0.6) in teal. Grey dots indicate non-significant values. The horizontal dashed line represents the p adj cutoff. (B) Venn diagrams describing upregulated and downregulated DEGs, common and unique to each comparison group (TET1 = TET1-high cancers, TET2 = TET2-high cancers, TET3 = TET3-high cancers). (C) gseGO enrichment results of DEGs (p adj < 0.05). A maximum of 7 significantly enriched and activated GO terms, including BP, CC, and MF, are shown for each TET group. (D) Ridge plot visualizing expression distributions of core enriched genes for GSEA-enriched and activated GO pathways in TET3-high versus TET3-low groups. Gradient color reflects the p adj value.

To investigate the biological pathways differentially enriched in the high- and low-TET groups, GSEA was conducted, showing a common enrichment in all TET-high samples in the activated biological process linked to RNA processing (GO:0006396, **Fig. 6C-D** and **Supplementary Fig. 5A-C, Supplementary Data 1**). Interestingly, uniquely for TET3-high and TET2-high samples, a further significant enrichment in two biological processes associated with protein-RNA complex (assembly and organization, GO:0022618 and GO:0071826, respectively) was also observed. The activation of the protein-RNA complex assembly process is a critical mechanism for cellular regulation and function. This process includes several key complexes, such as the ribosome, the spliceosome, the exosome complex, and the RNA-induced silencing complex (RISC). A KEGG pathway analysis notably highlighted the spliceosome as the only significant pathway activated in TET2 and TET3-high cancers, suggesting it as a possible explanation for the observed biological process (**Supplementary Fig. 5D** and **Supplementary Fig. 6, Supplementary Data 1**).

Overexpression of any TET was also linked to an increased abundance of nucleolus-associated genes (GO:0005730) and several small nucleolar RNAs (snoRNAs) and small Cajal body-specific RNAs (scaRNAs) (**Supplementary Fig. 10** and **Supplementary Fig. 5C, Supplementary Data 1**).

### TET3 expression aligns with RNA-processing signatures across breast cancer models

Given that our tissue and TCGA analyses showed the strongest cytoplasmic 5hmC and TET3 enrichment in triple-negative breast cancers, we focused functional validation on TNBC/basal models. To identify suitable *in vitro* systems, we first examined the baseline TET expression across the breast cancer cell lines analyzed in the DepMap resource. This revealed that TET2 and TET3 are expressed at substantially higher levels than TET1 (TET1 median = 0.48 ± 0.55 SD; TET2 median = 2.56 ± 0.80 SD; TET3 median = 2.69 ± 0.83 SD; all BH-adjusted p < 10⁻³³) (**Supplementary Fig. 8A**). Although TET2 and TET3 showed broadly similar expression, TET3 remained modestly but significantly higher (adjusted p = 1.80 × 10⁻³). Based on these profiles, we selected three representative TNBC cell lines spanning the diversity of TNBC subtypes and endogenous TET3 expression, corresponding roughly to the middle and upper quartiles of the distribution: MDA-MB-231 (basal-B), MDA-MB-468 (basal-A) and CAL-51 (basal-B) (**Supplementary Fig. 8A**). A marked reduction in TET3 mRNA expression, but not in TET2 and TET1 expression, was observed in all three cell lines following siTET3 treatment compared with scramble control, confirming knockdown specificity (paired two-tailed t-tests, TET3 CAL51: p = 0.0091; MDA-MB-231: p = 0.0004; MDA-MB-468: p = 0.010; TET1 and TET2, all p > 0.36) (**Supplementary Fig. 8B**). Knockdown was further confirmed at the protein level by immunofluorescence in MDA-MB-468 cells, which showed a significant reduction in TET3 signal in both the nucleus and the cytoplasm 48 h after siTET3 transfection compared with scrambled control (linear mixed-effects model on all cells, nucleus, cytoplasm and whole cell all p < 0.0001; Kruskal–Wallis/Dunn’s test on biological replicate medians, nucleus p = 0.050, whole cell p = 0.050, cytoplasm p = 0.127, n.s.) (**Supplementary Fig. 8C– D** and **Supplementary Table 7, Supplementary Data 1**).

We thus assessed whether TET3 depletion recapitulated the transcriptional patterns observed in TET3-high TCGA tumors by validation of a curated subset of TCGA/tissue-derived DEGs. Targets included (i) top-ranked tissue DEGs and (ii) genes with functional links to RNA processing/spliceosome, including both TET3-only and Shared (TET1/2/3) categories (**Supplementary Data 2**). Consistent with the tissue analysis, TET3 knockdown produced modest but directionally consistent decreases in a subset of RNA-processing and spliceosomal transcripts *in vitro* (**Supplementary Data 2**). Albeit with heterogeneous magnitude across individual genes and cell lines, the majority of genes with positive tissue log₂FC showed FC < 1 after knockdown, indicating a coherent trend towards downregulation (**Fig. 7A**). Significant decreases (mean fold-change siTET3/scramble, unadjusted p) were observed in MDA-MB-468 for two RNAs associated with RNA processing, SCARNA10 (0.38, p = 0.036) and SNORA53 (0.56, p = 0.033), for the spliceosomal protein CDC40 (0.61, p = 0.021), and for the transcriptional regulator HMX2 (0.38, p = 0.021); in CAL-51 for the spliceosomal components CDC40 (0.60, p = 0.005) and SF3B1 (0.85, p = 0.045); and in MDA-MB-231 for SNORA53 (1.53, p = 0.045; note this direction diverges from the tissue trend in this model) and the potassium channel KCNT1 (10.62, p = 0.050). Several additional RNA-processing/snoRNA candidates (SCARNA5, SCARNA7, SNORA23, SNORA73B, RN7SK, PRPF18, SMNDC1) showed the same direction (FC < 1) in one or more lines but did not reach significance at n = 3. Taken together, these data provide supporting evidence for a reduction in RNA-processing/spliceosome and snoRNA targets upon knockdown, most clearly in MDA-MB-468 and CAL-51 (**Fig. 7A** and **Supplementary Fig. 8E**).

**Fig. 7.**
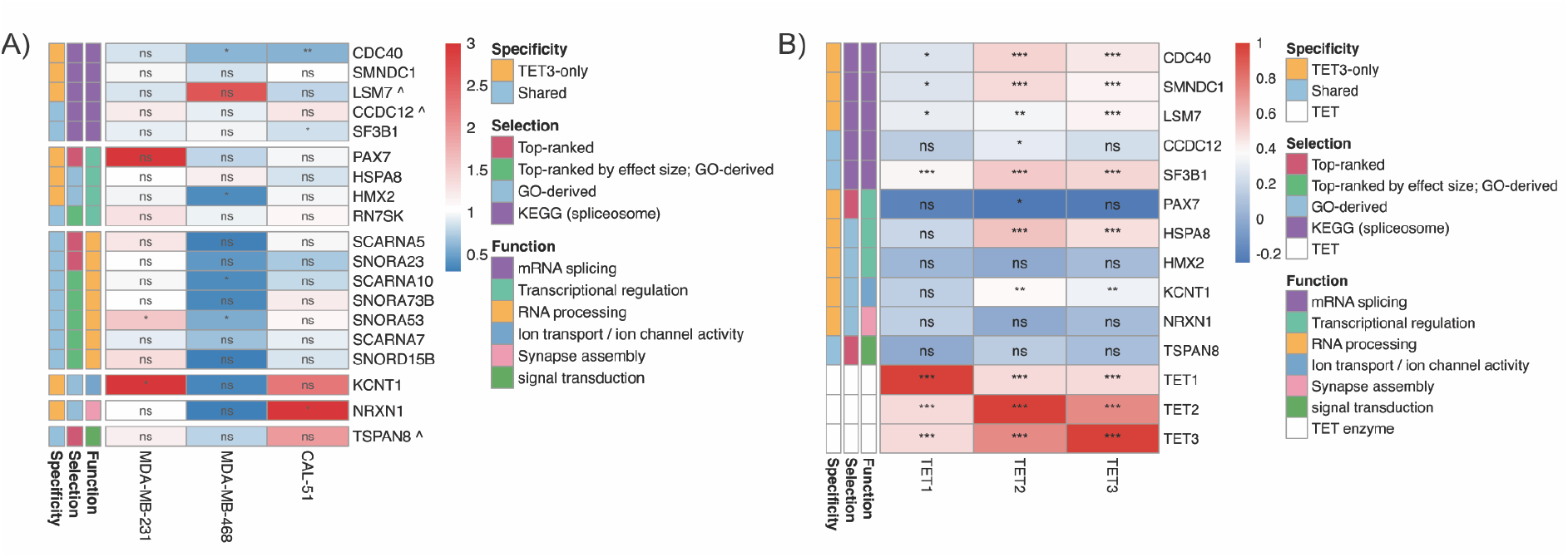
*TET3* regulates candidate target genes in breast cancer. (A) qPCR validation of candidate genes following *TET3* knockdown in three TNBC cell lines (MDA-MB-231, MDA-MB-468, CAL-51). Genes marked with a caret (^) correspond to transcripts that exhibited a negative log₂ fold change in the TCGA TET3-high versus TET3-low analysis. Heatmap shows fold-change (siTET3 / scramble) with color scale centered at 1 (blue = downregulation, red = upregulation). Genes are annotated by function (colored bars), TET3 specificity, and selection criteria. Statistical significance: * p < 0.05, ** p < 0.01, *** p < 0.001 (one-sample t-test on log₂FC vs. 0). Gaps separate functional categories. (B) Pearson correlations between TET genes (TET1, TET2, TET3; columns) and candidate target genes (rows) across breast cancer cell lines from DepMap. Color intensity represents correlation strength (blue = negative, red = positive). Numbers show correlation coefficients with significance stars (* p < 0.05, ** p < 0.01, *** p < 0.001; FDR-adjusted). Gene order matches panel A.

As the qPCR experiments were restricted to three breast cancer cell lines, to test whether the same TET3-associated subset of transcripts vary with TET3 expression across a broader set of cells, we analyzed baseline correlations across all 70 DepMap breast cancer cell lines. While this analysis is correlational rather than causal, it offers an orthogonal, independently derived dataset to contextualize and interpret the siTET3 findings. Although short RNAs (e.g., snoRNAs/scaRNAs) are not quantified in DepMap, two of the genes that were significantly reduced by siTET3 knockdown, CDC40 and SF3B1, also showed strong positive correlations with endogenous TET3 (CDC40: r = 0.447, padj = 1.04×10⁻⁴; SF3B1: r = 0.496, padj = 1.29×10⁻⁵) across the full DepMap breast panel. Beyond these top validated genes, several additional RNA-processing and spliceosome-related transcripts (e.g., RBM25, SMNDC1, LSM7, PPIH, SF3A2) also showed significant positive associations with TET3 in DepMap (**Fig. 7B** and **Supplementary Table 8, Supplementary Data 1**). Although these genes did not reach significance in all knockdown models, the consistency of their directionality across datasets further supports a broader link between TET family members and RNA-processing programs. As expected, TET2 and TET3 were strongly co-expressed across the panel, consistent with a partly shared TET-family transcriptional program. Importantly, subtype-stratified correlation analyses revealed that TET3–splicing associations were strongest in TNBC, particularly basal-B, mirroring both the tissue-level biology and our choice of TNBC models for functional validation (**Supplementary Fig. 9, Supplementary Data 1**).

Collectively, these tissue, TCGA, functional and independent cell-line datasets support TET3 as a leading candidate regulator of RNA-processing gene expression in breast cancer. However, as TET3 also hydroxymethylates DNA, the present data cannot yet distinguish a direct effect of TET3 on RNA hydroxymethylation from an indirect, DNA/chromatin-mediated route to the same transcriptional outcome.

## Discussion

In this study, we demonstrate the cytoplasmic localization of 5hmC in BC cells and tissues and provide direct evidence, through an RNase-sensitive, RNA-specific assay, for RNA hydroxymethylation in BC. Building on this primary finding, our analysis further supports TET3, and to a lesser extent TET2, as leading candidate regulators of transcripts involved in RNA processing, including mRNA modification and splicing, in this cancer type. The decrease in 5hmC levels has been considered a hallmark of cancer, as indicated by data obtained from various techniques including liquid chromatography–mass spectrometry (LC-MS/MS) for digested genomic DNA, 5hmC-specific sequencing, and immunoassays. Immunoassays allow a full morphological evaluation of the tissue, detection of 5hmC enrichment in all nucleic acids, and potential colocalization between TETs, 5hmC and subcellular structures such as Cajal bodies involved in RNA biogenesis and processing ^30^. The presence of scaRNAs in TET2 and TET3-high cancers from our analysis further is consistent with a potential involvement of TET proteins in Cajal bodies in BC.

Immunoassays can also distinguish between the presence of 5hmC in the nucleus, where DNA and several classes of RNA, notably pre-mRNA/unspliced mRNA, are located, and in the cytoplasm, where other RNA classes, including mature mRNA, tRNA, and rRNA, reside. While previous studies have quantified the nuclear signal of 5hmC in cancer tissues, cytoplasmic signals have been ignored. This is possibly due to heterogeneity in immunoassay protocols and omission of HCl pre-treatment to release nucleic acids from bound proteins, which could lead to underestimation of cytoplasmic signals ^25,31^.

Our study demonstrates the feasibility of detecting a strong and quantifiable cytoplasmic 5hmC signal in BC cell lines *in vitro* and even more prominently in human BC tissue samples. The weaker cytoplasmic signal observed in cell lines compared to human tissues aligns with previous findings reporting reduced levels of 5hmC measured in the DNA or in the nuclei *in vitro* ^16^, particularly in less differentiated cells ^25^.

Our study revealed that 5hmC is a feature of both total RNA and mRNA in BC cell lines, suggesting potential novel biological functions of RNA modifications in cancer, similar to 5mC modifications and of clinical significance ^32^. There are convincing reasons to consider 5hmC as a potential biomarker for BC and to investigate its prognostic significance in assessing disease prognosis and guiding treatment decisions (reviewed by ^33^). Our findings indicate the abundance of cytoplasmic 5hmC in our small cohort of ER-/PR-samples. This observation correlates with differences in TET2 and TET3 localization in our samples and in *TET3* expression analyzed from a paired normal-cancer TCGA dataset. Previous analyses using qPCR on small sample cohorts ^24,34^, and TCGA dataset analysis employing RNAseq data ^24^ support our findings, highlighting *TET3* as the most prominently expressed mRNA and its contribution to differences in 5hmC levels in BC.

While both oncogenic and tumor suppressor functions of *TET1*, *TET2*, and *TET3* in BC have been identified (reviewed by ^35^), prior studies have primarily focused on the interplay between TETs and DNA hydroxymethylation. However, recent research highlights the critical role of 5hmC and TETs in splicing regulation in brain, in peripheral lymphocytes, and in breast cancer ^36–39^. Our TCGA analysis supports these findings, suggesting a possible involvement of TET2 and TET3 in the spliceosome, and therefore their contribution to pre-mRNA processing. As TET3 was the only TET enzyme consistently upregulated in paired tumour–normal analyses, and because the candidate transcripts were derived specifically from the TET3-high TCGA signature, we focused our functional validation on TET3. Consistent with this, TET3 knockdown in three TNBC/basal cell lines led to reduced expression of selected RNA-processing and spliceosomal genes, including CDC40 and SF3B1, providing functional evidence that TET3 levels influence these transcripts. Moreover, baseline expression patterns across the DepMap breast cancer panel showed positive associations between endogenous TET3 and multiple RNA-processing genes, particularly within TNBC subtypes, offering independent support for a TET3-linked RNA-processing signature in breast cancer.

In summary, our study demonstrates that cytoplasmic 5hmC is a consistent feature of breast cancer, particularly in triple-negative disease, and correlates with increased cytoplasmic localization of TET2/TET3. Through analysis of patient tissues, TCGA datasets, functional perturbation experiments, and independent cell line data, we establish an association between TET3 expression and RNA-processing gene signatures, including spliceosomal components. While the mechanism, whether direct RNA modification or indirect transcriptional regulation, remains to be determined, our findings position TET3 as a candidate regulator of RNA-processing programmes in breast cancer and provide a basis for future studies investigating the functional consequences of TET-mediated modifications in RNA metabolism.

## Methods

### Cell lines

Human BC cell lines MDA-MB-231, BT474, SKBR3, MDA-MB-468, CAL-51 and MCF7 were procured from ATCC (American Type Culture Collection, ATCC, Teddington, UK) and cultured in Dulbecco’s Modified Eagle Medium (DMEM, Sigma, Gillingham, UK), supplemented with 10% Fetal Bovine Serum (Gibco, Loughborough, UK) and 1% L-glutamine (Gibco). Cells were maintained at 20% O2, 5% CO2, and 37°C. Regular mycoplasma testing was conducted. Cells were passaged at 70% confluency using 0.05% trypsin in EDTA.

### Immunofluorescence

Cells were seeded at a density of 50,000 cells per 1.8 cm² chamber in Nunc LabTek 4-chamber Permanox slides (ThermoFisher Scientific, Loughborough, UK) and allowed to attach overnight. Subsequently, cells were fixed with a 4% (w/v) methanol-free formaldehyde solution for 15 minutes (min) at room temperature (RT), and permeabilized with cold Triton X-100 (0.2% in PBS) for 15 mins.

For 5hmC detection only, the cells were incubated in 2N HCl for 30 mins and neutralized for 10 mins with 100 mM pH 8.5 Tris-HCl. All cells underwent a blocking step for 1 hour (hr) using 10% normal goat serum (ThermoFisher Scientific), followed by incubation with primary antibodies at +4 °C overnight. The primary antibodies used were anti-5-hmC (1:200 dilution, #39769, Active Motif), anti-Beta actin (1:500 dilution, # ab6276, Abcam), anti-EGFR (1:50 dilution, #2239, Cell Signaling Technology), and anti-TET3 antibody (1:100, #SAB2700682, Sigma-Aldrich). An isotype control antibody was used to ensure negative nonspecific staining of 5hmC (recombinant rabbit IgG, 1:100 dilution, #ab172730, Abcam). Incubation with each secondary antibody (goat anti-rabbit IgG, Alexa Fluor 488 and goat anti-mouse IgG, Alexa Fluor 568, both used at a 1:1000 dilution and from ThermoFisher Scientific) was performed for 1 hr at RT, followed by nuclear staining with DAPI for 15 mins at RT and mounting with ProLong Gold Antifade mountant (ThermoFisher Scientific).

### Immunofluorescence quantification

For each cell line, images of representative areas containing a minimum of 30 individual cells were acquired using an Aurox Clarity Light source pE4000 CoolLed confocal microscopy system (Aurox Ltd, Abingdon, UK) equipped with a 63X 1.4NA water objective and a pco.panda 4.2 sCMOS camera (Excelitas, Waltham, MA, USA). Z slices of 2.5 μm each were captured, resulting in a total of 8-12 slices depending on cell size. The signal corresponding to 5hmC was acquired in all images with identical exposure times.

Fluorescence intensity was quantified using a custom semi-automated macro in Fiji (ImageJ). For each image, the DAPI channel was projected using maximum intensity, while the 5hmC channel was projected using average intensity to preserve quantitative signal linearity. The actin or EGFR channel was projected using maximum intensity to aid whole-cell boundary delineation. Nuclear segmentation was performed automatically on the DAPI maximum projection using intensity thresholding and particle analysis, followed by manual review and correction of individual nuclear ROIs. Whole-cell boundaries were derived semi-manually from the actin or EGFR maximum projection, with automated pre-segmentation followed by per-cell user correction using a brush tool. Cytoplasmic regions were defined by subtracting the nuclear mask from the corresponding whole-cell mask. Mean intensity, area, and integrated density were measured in the 5hmC average projection for each compartment (nucleus, cytoplasm, whole cell) per cell.

Downstream analysis was conducted in R (v. 4.4.1) and all figures were produced using the package *ggplot2* (v. 3.5.2), unless otherwise specified. Nuclear and cytoplasmic measurements were paired per cell, and the following metrics were calculated: (i) raw mean fluorescence intensity per compartment; (ii) log₂ nuclear/cytoplasmic (N/C) intensity ratio (log₂(Mean_Intensity_Nucleus / Mean_Intensity_Cytoplasm)); and (iii) cytoplasmic fraction of total 5hmC signal, defined as the integrated density of the cytoplasm divided by the sum of nuclear and cytoplasmic integrated densities (F_cytoplasm = ID_cytoplasm / (ID_nucleus + ID_cytoplasm)), where integrated density = mean intensity × area (μm²).

Single suspected outliers were identified using Rosner’s generalised ESD test on log2-transformed compartment intensities after nuclear/cytoplasmic pairing (package *EnvStats*, v. 3.1.0; type I error = 0.05), and the corresponding paired cell was removed if an outlier was detected. Two cells were removed in total (one BT474 nucleus outlier, one MDA-MB-231 cytoplasm outlier), leaving 308 cells for analysis (BT474 n = 70, MCF7 n = 48, MDA-MB-231 n = 78, SKBR3 n = 112).

Within each cell line, differences in 5hmC intensity between nuclear and cytoplasmic compartments were assessed using a linear mixed-effects model (log2(intensity) ∼ Class + (1|Replicate), per cell line; packages *lme4* v. 1.1-37 and *lmerTest* v. 3.2-1), with biological replicate as a random intercept. The cytoplasmic fraction and log2 N/C ratio were each tested against fixed reference values (0.5 and 0, respectively) using one-sample linear mixed-effects models ((metric − reference) ∼ 1 + (1|Replicate), per cell line). Differences across cell lines were assessed using Kruskal-Wallis tests followed by Bonferroni-adjusted Dunn’s post-hoc comparisons on biological replicate medians (n = 3 per cell line; package *dunn.test* v. 1.3.6).

### Data acquisition and preprocessing

Gene expression data and cell line metadata were obtained from the Cancer Dependency Map (DepMap) Public 25Q3 release. Expression values were provided as log₂(TPM + 1) for human protein-coding genes. Model metadata included Oncotree lineage classifications, primary/metastatic status, sample collection site, and molecular subtype features. The dataset was filtered to retain only breast cancer cell lines (OncotreeLineage = "Breast"). For each cell line, the default entry for gene expression was selected, and redundant columns were removed to optimize data structure. Expression data were merged with clinical metadata, retaining cell line identifiers and subtype annotations. Breast cancer cell lines were classified into molecular subtypes based on the expression of established marker genes and proliferation signatures, following PAM50-like classification principles. The following marker genes were used for subtype classification: ESR1, PGR, GATA3, FOXA1, KRT8, KRT18; ERBB2, GRB7, KRT5, KRT14, KRT17, EGFR, CLDN3, CLDN4, CLDN7, VIM. A proliferation score was calculated as the mean z-score of nine proliferation-associated genes: MKI67, UBE2C, BIRC5, NDC80, CCNB1, CDC20, PTTG1, RRM2, and MYBL2. The classification proceeded as follows: 1) Normal-like/Immortalized. Cell lines with known non-malignant origins (HMEL, HBL-100, MCF 10A, MCF10A) were classified first and excluded from cancer subtype analysis, as these represent immortalized normal mammary epithelial cells rather than tumor-derived lines. 2) HER2+. Cell lines with high ERBB2 expression (z-score ≥ 1) were classified as HER2+, regardless of ER status. By classifying HER2+ cases before Luminal cases, we ensure that HER2+/ER+ cell lines are grouped with HER2-enriched rather than Luminal subtypes, consistent with PAM50 molecular classification. 3) Luminal (ER+). High ESR1 expression (z-score ≥ 0.5), or high expression of both luminal transcription factors GATA3 and FOXA1 (z-scores ≥ 0.5). 4) TNBC Basal-B/Claudin-low. High VIM expression (z-score ≥ 0.5), or low expression of ≥2 claudin genes (CLDN3, CLDN4, CLDN7; z-scores ≤ - 0.3). 5) TNBC Basal-A. Remaining triple-negative cell lines were classified as Basal-A if they expressed any basal/myoepithelial marker at high levels: KRT5, KRT14, KRT17, or EGFR (z-score ≥ 1). 6) Unclassified. Cell lines not meeting any of the above criteria were labeled as "Unclassified." These represent cell lines with ambiguous or intermediate marker expression patterns that do not fit established molecular subtypes.

### Cell line selection and gene selection for validation of TCGA-derived DEGs

To select biologically relevant in-vitro models for functional validation, we first focused on triple-negative/basal breast cancer cell lines, as this subgroup showed the strongest cytoplasmic 5hmC and TET3 signals in our tissue analyses. Baseline expression of TET1, TET2, and TET3 across breast cancer cell lines was next assessed using transcriptomic data from the DepMap database. Across 70 breast cancer models, TNBC/basal lines exhibited moderate to high TET3 expression, making them suitable candidates for TET3 knockdown studies. Based on this, MDA-MB-231 (TNBC basal-B), MDA-MB-468 (TNBC basal-A), and CAL-51 (basal-B) were selected to capture the natural variation in TET3 expression and TNBC/basal subtype diversity.

### siRNA Transfection of Cell Lines (TET3 Knockdown)

MDA-MB-231, CAL-51, and MDA-MB-468 were transfected with siRNAs targeting TET3 using Lipofectamine RNAiMAX (Thermo Fisher Scientific) following the manufacturer’s instructions with minor adaptations. Briefly, cells were seeded in 6-well plates in antibiotic-free full growth medium to achieve ∼50% confluence the following day. Cells were transfected with ON-TARGET plus SMART pool of TET3-targeting siRNAs or ON-TARGET non-targeting scrambled siRNA (Horizon Discovery, Cambridge, UK) at a final concentration of 40 nM. Target sequences of siRNA are provided in **Supplementary Table 6, Supplementary Data 1**.

Forty-eight hours post-transfection, cells were harvested for RNA extraction, as described below. All experiments were performed in three biological replicates. Knockdown efficiency and specificity were assessed by comparing TET1, TET2, and TET3 expression between scrambled and siTET3 conditions using paired t-tests for each cell line and gene combination. To confirm TET3 knockdown at the protein level, MDA-MB-468 cells were seeded onto Nunc LabTek chamber slides and transfected with TET3-targeting or scrambled siRNA as described above. Forty-eight hours post-transfection, cells were fixed, permeabilized and stained as described in the immunofluorescence method section, using anti-TET3 and anti-EGFR (the latter to delineate whole-cell boundaries) antibodies alongside DAPI. Nuclear, cytoplasmic and whole-cell TET3 intensities were quantified per cell as described above, from n = 3 biological replicates per condition (scrambled: 136 cells; siTET3: 147 cells). Differences in TET3 intensity between scrambled and siTET3 conditions were assessed within each compartment using a linear mixed-effects model (log2(intensity) ∼ Condition + (1|Replicate); packages *lme4*v. 1.1-37 and *lmerTest* v. 3.2-1), with biological replicate as a random intercept, and cross-checked using a Kruskal-Wallis test with Dunn’s post-hoc (Bonferroni-adjusted) on biological replicate medians (package *dunn.test* v. 1.3.6).

### Selection of DEG targets for validation

To validate candidate targets after TET3 knockdown in these cell lines, DEGs previously identified in breast cancer samples and stratified by TET3 expression (high vs. low) were ranked according to adjusted p value and filtered using biologically relevant criteria. For DEGs uniquely altered in TET3-high cancers ("TET3-only DEGs"), a threshold of log₂ fold change > 3 or < -3 and adjusted p value < 1.2×10⁻³ was applied to retain high-confidence, large-effect genes specific to TET3. Additionally, uniquely TET3-regulated DEGs identified through the GO enrichment analysis ("GO-derived DEGs") were included to capture functionally relevant pathways. A more stringent threshold (log₂ fold change > 2 or < –2 with significant adjusted p-values) was applied to ensure selection of robust, TET3-specific alterations, including those linked to enriched pathways. Finally, from the TET3-unique spliceosome-associated DEGs ("KEGG spliceosome"), we selected an initial subset of genes with significant adjusted p-values for validation. These genes represented different functional nodes within the spliceosome and showed consistent expression shifts. To distinguish these TET3-specific alterations from broader TET-family transcriptional responses, we next identified "common DEGs", defined as genes also differentially expressed in TET1-high or TET2-high cancers. For these set of candidates, the criteria for selection was log₂ fold change > 7 or < 0 and adjusted p value < 5×10⁻¹¹. This threshold was chosen to prioritize the most robust, high-confidence events, representing core transcriptional effects consistently associated with TET-family activity. Targets with undetermined or extremely low baseline signal (Ct ≳ 30) in the scramble control were excluded from validation analysis. The full list of validated genes and primer sequences is provided in **Supplementary Data 2**.

For downstream target genes, one-sample t-tests were performed on log₂(fold-change) values to test the null hypothesis that mean log₂(FC) = 0 (i.e., no change from scramble control). P-values were calculated for each gene × cell line combination.

### Correlation analysis in DepMap breast cancer cell lines

To provide an independent, orthogonal dataset for evaluating whether candidate DEGs co-vary with TET expression, we analyzed baseline RNA-seq (log2(TPM+1)) expression values across DepMap breast cancer lines. Pairwise Pearson correlation coefficients were calculated between each TET gene and each target gene across all breast cancer cell lines using the cor.test() function in R. Confidence intervals (95%) were computed for each correlation. P-values were adjusted for multiple testing using the Benjamini-Hochberg FDR method. Correlations with FDR-adjusted p < 0.05 were considered statistically significant. Correlation matrices were visualized using the *pheatmap* package in R.

### RNA extraction and Poly(A) mRNA Enrichment

Total RNA was isolated using Zymo Quick-RNA Miniprep Kit (Zymo Research, Irvine, California, USA), according to the manufacturer’s protocol and quantitated using a NanoDrop One spectrophotometer (ThermoFisher Scientific). The integrity of total RNA, measured as a RIN number, was assessed with the Bioanalyzer RNA 6000 Nano assay on the Bioanalyzer system (Agilent, Santa Clara, California, USA). According to the manufacturer’s recommendations for PolyA mRNA enrichment, RNA with a RIN score > 7 was deemed appropriate. Enrichment of the polyadenylated (poly(A)) fraction of isolated total RNA was performed using the Dynabeads™ mRNA Purification Kit (ThermoFisher Scientific), following the manufacturer’s instructions. Briefly, oligo(dT) magnetic beads were employed to selectively capture the mRNA fraction from the total RNA pool (approximately 1% yield of mRNA from total RNA is expected). The captured poly(A) mRNA was then eluted from the beads in RNAse-free water and the concentration was measured using a NanoDrop One spectrophotometer (ThermoFisher Scientific). To further reduce residual rRNA contamination, the eluted mRNA was subjected to a second round of poly(A) capture using the same Dynabeads™ Oligo(dT) beads, following the manufacturer’s recommended "double isolation" protocol.

### Real-time quantitative polymerase chain reaction (qPCR)

First-strand cDNA synthesis from total RNA was performed using the RevertAid First Strand cDNA Synthesis Kit (ThermoFisher Scientific) using random hexamer primers and according to the manufacturer’s instructions.

The expression of TET1, TET2, and TET3 and the housekeeping gene HPRT1 (Hypoxanthine-Guanine Phosphoribosyltransferase 1) was determined using the StepOne Plus Real-Time PCR System software (ThermoFisher Scientific). Primers were verified for specificity via a BLAST search against the GenBank database (National Institutes of Health, Bethesda, MD, USA) or designed using the Primer3 design tool^40^. Primer sequences are reported in **Supplementary Table 6, Supplementary Data 1**. To determine the optimized annealing temperature for all primers, gradient PCR reactions were executed on cDNA samples derived from the BT474 cell line, employing Taq DNA Polymerase (Qiagen). The PCR conditions were as follows: initial denaturation at 94°C for 5 minutes; 34 cycles of denaturation at 94°C for 30 seconds, annealing at 54-65°C for 40 seconds, and elongation at 72°C for 30 seconds; followed by a final elongation at 72°C for 10 minutes. Electrophoresis on a 1.5% agarose gel with a DNA Hyperladder 100bp (Bioline, London, UK) confirmed the presence of a single specific PCR band. To assess primer efficiency and determine the appropriate concentrations of both primers and cDNA for subsequent reactions, qPCR experiments were conducted at the optimized annealing temperature. Serially diluted concentrations of cDNA samples (20, 2, 0.2, 0.02 ng/μl) were used. The standard curve and melting curve analyses were performed using the StepOne Plus Real-Time PCR System software (ThermoFisher Scientific). The qPCR conditions comprised 40 cycles of denaturation at 95°C for 3 seconds and annealing at 63°C for 30 seconds. Primer efficiency slopes were compared to ensure uniformity, and melting curve analysis was conducted to verify the absence of multiple peaks indicative of primer aspecificity or primer dimer formation. Gene expression was quantified using the ΔΔCt method, using TET1 in MDA-MB-231 as the baseline. Final ΔΔCt values represent fold change in gene expression relative to the baseline, normalized to the reference gene. A one-way ANOVA was performed to test the difference of the signal in each cell line, followed by a Tukey’s HSD post-hoc test. A p-value equal to or less than 0.05 was considered statistically significant and, in the figures, indicated with an asterisk (*p<0.05). Increasing statistical significance is indicated with more asterisk: *p<0.05, **p<0.01, ***p<0.001, ****p<0.0001.

### Detection of 5hmC in RNA

A modified MethylFlash™ 5-mC RNA Methylation ELISA Easy Kit (Epigentek, Farmingdale, NY, USA) was used to quantify 5hmC content in RNA. The provided 5mC antibody was substituted with a 5hmC antibody (A-1018, Epigentek). A 1:100 dilution in H2O of a hydroxymethylated polynucleotide containing 20% of 5hmC from a commercial kit (MethylFlash™ Hydroxymethylated DNA Quantification Kit, Epigentek) was used as a positive control and loaded in equal amounts to the RNA samples. Otherwise, the manufacturer’s instructions were followed. The input of both mRNA and total RNA samples was 300 ng. The fluorescence intensity was measured at 530ex/590em nm on an Infinite M1000 microplate reader (Tecan, Männedorf, Switzerland).

### Assay validation and specificity controls

To ensure that the detected 5hmC signal originated from RNA rather than contaminating DNA or non-specific binding, we performed a series of validation experiments (**Fig. 2**). Bioanalyzer electropherograms of RNA samples post-extraction were obtained to confirm RNA integrity and the absence of detectable genomic DNA contamination. An RNA dilution series was performed using DNase-treated total RNA at four input amounts (300 ng, 150 ng, 75 ng, 0 ng) to assess the dose-dependence of the signal. A subset of RNA samples was treated with RNase I prior to the assay to confirm RNA-dependence of the signal. Finally, serial dilutions of 5hmC-DNA positive controls were run on the RNA-capture ELISA plate to assess cross-reactivity with DNA.

### Human tissue samples

#### Colon tissue

A cohort of 20 formalin-fixed paraffin-embedded (FFPE) tissue sections was obtained from 7 patients affected by inflammatory bowel disease who have undergone colectomy for colitis-associated cancer (CAC) at St. Mark’s Hospital, Harrow, UK (ethical approval from London North West NHS Trust Research ref: RD/16/084). The age of the patients ranged from 30 to 60, and male to female ratio was 5:2. The cohort comprised 12 non-malignant (quiescent) sections of colon tissue and, from a subset of 5 patients, 8 matched colitis-associated cancer tissues were available.

#### Breast tissue

A cohort of 29 patients, including 21 pairs of invasive ductal carcinoma (IDC) from post-menopausal patients between 40 to 68 with matching sections containing phenotypically normal breast from an adjacent area of the same patient was collected from Imperial College Tissue Bank (ICHTB). ICHTB is supported by the National Institute for Health Research (NIHR) Biomedical Research Centre based at Imperial College Healthcare NHS Trust and Imperial College London, and it is approved by Wales REC to release human material for research (17/WA/0161). This work was performed following the ethical principles in the Declaration of Helsinki. See **Supplementary Table 1** for clinical information.

To confirm the normal histology, hematoxylin and eosin (H&E) staining was performed on the first section of each sample. Each section was cut at 5μm thickness, hydrated in decreasing concentration of ethanol, and stained using Gills Hematoxylin and 1% Eosin, as per standard protocol.

### Immunohistochemistry (IHC)

Five-micron thick FFPE tissue sections were dewaxed, rehydrated and subjected to boiling in 10 mM sodium citrate buffer solution, pH 6.0 (Sigma-Aldrich, UK) for 20 min. For the detection of 5hmC only, the sections were incubated with hydrochloric acid (HCl, 2M) for 15 min at RT, then neutralized for 10 min using 300ul 100mM pH 8.5 Tris-HCl. All sections were subject to incubation with 10% goat serum (ThermoFisher Scientific, UK) for 10 min. Endogenous peroxidase activity was further blocked with 3% hydrogen peroxide solution for 10 min. Sections were then incubated for at 4°C overnight with the primary antibodies: 5hmC (Active motif, #39769, 1:5000 dilution in goat serum), 5mC (Cell Signalling Technology, USA, #CST28692, 1:1000), TET1 (Sigma-Aldrich, #HPA019032, 1:50), TET2 (Abcam, UK, #ab94580, 1:100), followed by PBS washes and incubation for 40 min at RT in biotin-conjugated goat anti-rabbit IgG (Agilent Technologies, UK, #0432, 1:1000 dilution in goat serum). Sections were then incubated in streptavidin-conjugated HRP (Cell Signalling Technologies, 1:200 in goat serum) for 40 min at room temperature. The color was developed with a DAB Peroxidase (HRP) Substrate Kit (Vector Laboratories, UK) according to the manufacturers’ recommendations and counterstained with hematoxylin, before dehydration through alcohol, clearing in xylene and mounting. For each tissue, the first serial section was separately stained with hematoxylin and eosin (H&E) to determine tissue morphology and distinguish cancer areas.

### Immunohistochemistry quantification

The images of each stained section were first digitally acquired on a Nanozoomer SQ slide scanner (Hamamatsu Photonics, Welwyn Garden City, UK). Tissues stained for 5hmC, 5mC and TET enzymes underwent a manual scoring by two independent observers and quantification with either the commercially available Visiopharm software (Visiopharm, Horsham, Denmark) or the open source QuPath software (41). The regions of interest (ROIs) were selected from the respective H&E images and quantification was performed blind. A minimum of 100 cells per region and a minimum of 3 regions were selected for each manual scoring unless a section of normal tissue was characterized by a minimal amount of epithelium. In that case, all quantifiable areas were evaluated. Since the samples were collected and processed at different times, to minimize the effect of differences in the quality and of the signal amongst the individual tissues, an internal normalization was achieved by identifying areas of negative stain (either DAB-negative nuclei, cytoplasm, or areas with minimal cellular component, such as connective tissue) to compare and subtract any background. For manual scoring, any signal from the cytoplasm, regardless of its strength, that was visibly stronger than the negative control was considered positive. The percentage of cells containing positive cytoplasm was determined by counting the number of positive cells against the total number of cells in the area.

Similarly, for the two automatic software methods, a generic threshold was initially generated to detect cytoplasmic signals based on color detection parameters established by each software. This threshold was then adjusted for each tissue using a training region of interest (ROI) containing a mixture of positive and negative cells, with the negative control as reference. The manual scoring obtained from manual ROIs, which had been agreed upon by two independent observers, was used to further refine the settings for both software methods within the training ROI. The classifier was subsequently applied to all ROIs to determine the percentage of cells with positive cytoplasm in each sample.

In the case of QuPath software, the image type was set to "BRIGHTFIELD_H_DAB". Deconvolution stain thresholds were determined using the aforementioned training ROIs, which included a negative control and regions with both positive and negative signals. Cell number, positive nuclei, and positive cytoplasm detection were conducted within each annotation using the PositiveCellDetection plugin, utilizing signals detected via Optical Density Sum (for cell number), Nucleus: DAB OD mean (for positive nuclei), and Cytoplasm: DAB OD mean (for positive cytoplasm).

The agreement between the two automatic methods was assessed on the cohort of breast tissues evaluated for 5hmC using a Bland-Altman plot (**Supplementary Fig. 7, Supplementary Data 1**), which indicated a bias of 7.883 +/- 22.19.

Statistical analysis was conducted in R (v. 4.4.1) and all figures were produced using the package ggplot2 (v. 3.5.1), unless otherwise specified. A p-value equal to or less than 0.05 was considered statistically significant and indicated with an asterisk (*p<0.05). Increasing statistical significance is indicated with more asterisk: *p<0.05, **p<0.01, ***p<0.001, ****p<0.0001.

### Analysis of TCGA data and identification of differentially expressed genes

Clinical data and gene expression data of BC female patients from the GDC TCGA Breast Cancer (BRCA) cohort were obtained from The Cancer Genome Atlas Program (TCGA) (https://www.cancer.gov/ccg/research/genome-sequencing/tcga) using cBioportal (http://cbioportal.org) and UCSC Xena (http://xena.ucsc.edu/), respectively. A total of 823 primary tumor samples and 102 non-tumor samples (defined as solid tissue normal) of invasive ductal carcinoma were selected, for which gene expression data and receptor status were available. The R package UCSCXenaTools (v. 1.4.8) ^41^ was utilized to extract gene expression data from the GDC TCGA Breast Cancer (BRCA) cohort (datasets "gene expression RNAseq - HTSeq - FPKM-UQ" and "gene expression RNAseq - HTSeq – Counts") from the GDC Xena Hub. Expression levels of ER, PR, and HER2 were determined based on the information corresponding to the primary tumors, which was downloaded from the dataset "TCGA-BRCA.GDC_phenotype" and integrated from cBioPortal, whenever possible for missing data. To determine if there were any statistically significant differences between the means of TET expression in multiple groups, a one-way ANOVA was performed, followed by a Tukey’s HSD post-hoc test. Paired t-test was conducted on patients for whom both primary tumor and matching normal tissue samples were available (101 patients). A p-value equal to or less than 0.05 was considered statistically significant and, in the figures, indicated with an asterisk (*p<0.05). Increasing statistical significance is indicated with more asterisk: *p<0.05, **p<0.01, ***p<0.001.

The differential expression analysis was performed using the R package DESeq2 (v.1.44.0) ^42^ using the raw count data. Only genes with a count of at least 10 in at least 10% of the samples were included in the analysis to ensure sufficient read coverage. Samples were stratified based on the normalized expression levels of the three TET mRNAs (TET1, TET2, and TET3) in cancer tissues relative to their matched normal tissues. "TET-high" cancer samples were defined as those in which TET mRNA expression was higher in cancer tissues compared to the matched normal tissues. Conversely, "TET-low" samples indicated higher TET mRNA expression in normal tissues than in cancer tissues. The variance stabilizing transformation (VST) was applied to the normalized counts for visualization and clustering purposes. Genes were considered significantly differentially expressed if p adj < 0.05 (Benjamini-Hochberg method). Volcano plots and heatmap were constructed using the R packages ggplot2 (v. 3.5.1) and pheatmap (v. 1.0.12), respectively. Venn diagrams were constructed using BioVenn ^43^

### Functional enrichment analysis

To investigate the biological pathways differentially enriched between TET-high and TET-low samples, Gene Set Enrichment Analysis (GSEA) was conducted using the R package clusterProfiler (v. 4.12.0) ^44^. This analysis was performed on the ranked list of differentially expressed genes to identify enriched Gene Ontology (GO) terms (BP, Biologic Process; CC, cellular component; MF, molecular function) and Kyoto Encyclopedia of Genes and Genomes (KEGG) pathways, using the functions gseGO and gseKEGG, respectively. The minimum gene set size (minGSSize) was set at 25 for GO terms to ensure meaningful biological interpretations, and at 10 for KEGG terms to capture relevant pathways with sufficient gene representation. Significantly enriched pathways were defined as pathways with p adj < 0.05 (Benjamini-Hochberg method).

## List of abbreviations

5mC: 5-methylcytosine
5hmC: 5-hydroxymethylcytosine
2OG: 2-oxoglutarate
BC: Breast cancer
CTNNB1: Catenin Beta 1
ER: Estrogen receptor
HER2: Human epidermal growth factor receptor 2
IDC: Invasive ductal carcinoma
IDH: Isocitrate dehydrogenase
IHC: Immunohistochemistry
LC-MS/MS: Liquid chromatography–mass spectrometry/mass spectrometry
PR: Progesterone receptor
qPCR: Quantitative polymerase chain reaction
TCGA: The Cancer Genome Atlas
TET: Ten-eleven translocation

## Declarations

### Ethics approval and consent to participate

Human colon tissue was collected at St. Mark’s Hospital, Harrow, UK (ethical approval from London North West NHS Trust Research ref: RD/16/084). Human breast tissue was collected from Imperial College Tissue Bank (ICHTB, ethical approval from Wales REC ref:17/WA/0161). This work was performed following the ethical principles in the Declaration of Helsinki with informed consent obtained from all subjects.

### Consent for publication

Not Applicable.

### Data availability

The dataset supporting the conclusions of this article is included within the article and its additional files.

### Competing interests

Three authors, JS, BC and NM, serve as editors for Oncogene, part of Springer Nature Group. The editorial roles did not influence the research presented here. From 2022 - present JS has been Editor-in-Chief of Oncogene and has sat on SABs/advisory boards for Lilly, Agenus, Celltrion, Greenmantle, vTv Therapeutics, APIM, Onconox, IO Labs, Clinical Ink, Zephyr AI, Benevolent AI, Biozen, Pear Bio, Sable Bio and Linkgevity. He has consulted with Lansdowne partners and Vitruvian. Since 2024 he has been a board member for Graviton Bioscience BV, Etira Therapeutics and Portage, and previously chaired the board of BB Biotech Healthcare Trust PLC. None of these interests are relevant here. The other authors declare no competing interests.

### Author contributions

JW performed data curation, formal analysis, investigation, and methodology. HY, KG, SS, FP, LEE, KM, PC, EH performed the investigation and formal analysis. LDDB, SO and NM performed supervision and validation analysis. JS conceptualized the study and provided resources and funding acquisition. BC conceptualized the study, performed data curation, formal analysis, investigation, and methodology, and wrote the original draft. All authors reviewed and edited the final draft of the submitted version.

## Acknowledgments

The work is funded by Action Against Cancer (grants P71130, JW; P66814, LDBB; P71728, NM; P62625, BC). The funders had no role in study design, data collection and analysis, decision to publish, or preparation of the manuscript.

We acknowledge the support of the NIHR and Imperial BRC and ECMC. We would like to thank the late Hilary Craft OBE, the visionary founder of Action Against Cancer, whose unwavering support made this research possible. Finally, we would like to thank the late Prof Sami Shousha (Imperial College London) for histopathology advice.

## Supplementary Data 1 – content

**Supplementary Figure 1.** Hydroxymethylation in the total RNA and mRNA of BC cells, with technical positive control.

**Supplementary Figure 2.** 5mC is predominantly a nuclear feature of BC tissue.

**Supplementary Figure 3.** Levels of 5hmC are higher in colon cancer tissue.

**Supplementary Figure 4.** TET3 expression differences in the BCD cohort.

**Supplementary Figure 5.** Gene Set Enrichment Analysis associated with each TET in the BCD cohort.

**Supplementary Figure 6.** Spliceosome pathway visualization.

**Supplementary Figure 7.** Comparison of IHC quantification methods.

**Supplementary Figure 8.** TET expression in breast cancer and validation of TET3-regulated genes.

**Supplementary Figure 9.** Pearson correlation coefficients between TET1, TET2, TET3 and candidate target genes within breast cancer molecular subtypes.

**Supplementary Figure 10.** TET-family-associated nucleolar and small nucleolar/Cajal body RNA (snoRNA/scaRNA) gene signature in the BCD cohort.

**Supplementary Table 1.** Clinical information for BC samples analyzed in the study.

**Supplementary Table 2.** Statistical analyses of 5hmC fluorescence intensity in nuclear and cytoplasmic compartments of four BC cell lines.

**Supplementary Table 3.** Results of one-way ANOVA and Tukey’s post-hoc test for TET mRNA expression levels among cell lines.

**Supplementary Table 4.** P-values of paired and unpaired tests of 5hmC expression from IHC in normal-cancer pairs.

**Supplementary Table 5.** Statistical analyses of TET mRNA expression patterns in BC samples derived from TCGA.

**Supplementary Table 6.** qPCR and siRNA sequences used in this study.

**Supplementary Table 7.** Statistical analyses of TET3 fluorescence intensity after knockdown.

**Supplementary Table 8.** Correlation of selected genes with TET family expression in DepMap.

## Supplementary Data 2

Full list of validated DEG targets, and selection criteria for functional validation.

## Notes

### Competing Interest Statement

The authors have declared no competing interest.

### Summary of Updates

Figure 6 updated to correct an unintentionally hidden label in the Venn diagram. Title of Supplementary file 1 updated to reflect the manuscript.

## References

1 Holm, K. et al. Molecular subtypes of breast cancer are associated with characteristic DNA methylation patterns. Breast Cancer Res 12, R36 (2010). 10.1186/bcr2590

2 Bediaga, N. G. et al. DNA methylation epigenotypes in breast cancer molecular subtypes. Breast Cancer Res 12, R77 (2010). 10.1186/bcr2721

3 Sadzeviciene, I. et al. Analysis of Intrinsic Breast Cancer Subtypes: The Clinical Utility of Epigenetic Biomarkers and TP53 Mutation Status in Triple-Negative Cases. Int J Mol Sci 23 (2022). 10.3390/ijms232315429

4 Schröder, R. et al. The epigenetics of breast cancer - Opportunities for diagnostics, risk stratification and therapy. Epigenetics 17, 612–624 (2022). 10.1080/15592294.2021.1940644

5 Jin, S. G. et al. 5-Hydroxymethylcytosine is strongly depleted in human cancers but its levels do not correlate with IDH1 mutations. Cancer Res 71, 7360–7365 (2011). 10.1158/0008-5472.Can-11-2023

6 Li, X., Liu, Y., Salz, T., Hansen, K. D. & Feinberg, A. Whole-genome analysis of the methylome and hydroxymethylome in normal and malignant lung and liver. Genome Res 26, 1730–1741 (2016). 10.1101/gr.211854.116

7 Kudo, Y. et al. Loss of 5-hydroxymethylcytosine is accompanied with malignant cellular transformation. Cancer Sci 103, 670–676 (2012). 10.1111/j.1349-7006.2012.02213.x

8 Fujikura, K. et al. Downregulation of 5-hydroxymethylcytosine is an early event in pancreatic tumorigenesis. J Pathol 254, 279–288 (2021). 10.1002/path.5682

9 Sjöström, M. et al. The 5-Hydroxymethylcytosine Landscape of Prostate Cancer. Cancer Res 82, 3888–3902 (2022). 10.1158/0008-5472.Can-22-1123

10 Yang, Q., et al. Decreased 5-hydroxymethylcytosine (5-hmC) is an independent poor prognostic factor in gastric cancer patients. J Biomed Nanotechnol 9, 1607–1616 (2013). 10.1166/jbn.2013.1713

11 Gambichler, T., Sand, M. & Skrygan, M. Loss of 5-hydroxymethylcytosine and ten-eleven translocation 2 protein expression in malignant melanoma. Melanoma Res 23, 218–220 (2013). 10.1097/CMR.0b013e32835f9bd4

12 Lian, C. G. et al. Loss of 5-hydroxymethylcytosine is an epigenetic hallmark of melanoma. Cell 150, 1135–1146 (2012). 10.1016/j.cell.2012.07.033

13 Orr, B. A., Haffner, M. C., Nelson, W. G., Yegnasubramanian, S. & Eberhart, C. G. Decreased 5-hydroxymethylcytosine is associated with neural progenitor phenotype in normal brain and shorter survival in malignant glioma. PLoS One 7, e41036 (2012). 10.1371/journal.pone.0041036

14 Munari, E. et al. Global 5-Hydroxymethylcytosine Levels Are Profoundly Reduced in Multiple Genitourinary Malignancies. PLoS One 11, e0146302 (2016). 10.1371/journal.pone.0146302

15 Li, W. & Liu, M. Distribution of 5-hydroxymethylcytosine in different human tissues. J Nucleic Acids 2011, 870726 (2011). 10.4061/2011/870726

16 Nestor, C. E. et al. Tissue type is a major modifier of the 5-hydroxymethylcytosine content of human genes. Genome Res 22, 467–477 (2012). 10.1101/gr.126417.111

17 Li, M. et al. 5-methylcytosine RNA methyltransferases and their potential roles in cancer. J Transl Med 20, 214 (2022). 10.1186/s12967-022-03427-2

18 Yi, J. et al. Overexpression of NSUN2 by DNA hypomethylation is associated with metastatic progression in human breast cancer. Oncotarget 8, 20751–20765 (2017). 10.18632/oncotarget.10612

19 Lan, J. et al. Functional role of Tet-mediated RNA hydroxymethylcytosine in mouse ES cells and during differentiation. Nat Commun 11, 4956 (2020). 10.1038/s41467-020-18729-6

20 Xia, M. et al. The Tet2-Upf1 complex modulates mRNA stability under stress conditions. Front Genet 14, 1158954 (2023). 10.3389/fgene.2023.1158954

21 Wu, R. et al. NSUN5/TET2-directed chromatin-associated RNA modification of 5-methylcytosine to 5-hydroxymethylcytosine governs glioma immune evasion. Proc Natl Acad Sci U S A 121, e2321611121 (2024). 10.1073/pnas.2321611121

22 Huang, W. et al. Formation and determination of the oxidation products of 5-methylcytosine in RNA. Chem Sci 7, 5495–5502 (2016). 10.1039/c6sc01589a

23 Fu, L. et al. Tet-mediated formation of 5-hydroxymethylcytosine in RNA. J Am Chem Soc 136, 11582–11585 (2014). 10.1021/ja505305z

24 Yang, H. et al. Tumor development is associated with decrease of TET gene expression and 5-methylcytosine hydroxylation. Oncogene 32, 663–669 (2013). 10.1038/onc.2012.67

25 Haffner, M. C. et al. Global 5-hydroxymethylcytosine content is significantly reduced in tissue stem/progenitor cell compartments and in human cancers. Oncotarget 2, 627–637 (2011). 10.18632/oncotarget.316

26 Broome, R. et al. TET2 is a component of the estrogen receptor complex and controls 5mC to 5hmC conversion at estrogen receptor cis-regulatory regions. Cell Rep 34, 108776 (2021). 10.1016/j.celrep.2021.108776

27 Tsai, K. W. et al. Reduction of global 5-hydroxymethylcytosine is a poor prognostic factor in breast cancer patients, especially for an ER/PR-negative subtype. Breast Cancer Res Treat 153, 219–234 (2015). 10.1007/s10549-015-3525-x

28 Kim, M. R., Wu, M. J., Zhang, Y., Yang, J. Y. & Chang, C. J. TET2 directs mammary luminal cell differentiation and endocrine response. Nat Commun 11, 4642 (2020). 10.1038/s41467-020-18129-w

29 Zhang, Z. et al. Values of 5mC, 5hmC, and TET2 for identifying the presence and progression of breast precancerous lesion. J Clin Lab Anal 34, e23162 (2020). 10.1002/jcla.23162

30 El Osmani, N., et al. Vitamin & enhances co-localization of novel TET1 nuclear bodies with both Cajal and PML bodies in colorectal cancer cells. Epigenetics 19, 2337142 (2024). 10.1080/15592294.2024.2337142

31 Zhong, S., Li, Z., Jiang, T., Li, X. & Wang, H. Immunofluorescence Imaging Strategy for Evaluation of the Accessibility of DNA 5-Hydroxymethylcytosine in Chromatins. Anal Chem 89, 5702–5706 (2017). 10.1021/acs.analchem.7b01428

32 Zhang, Z., Zeng, C. & Zhang, W. Considerations before the application of 5-hydroxymethylation levels of long non-coding RNAs for non-invasive cancer diagnosis. Extracell Vesicles Circ Nucl Acids 3, 10–13 (2022). 10.20517/evcna.2021.22

33 Xu, T. & Gao, H. Hydroxymethylation and tumors: can 5-hydroxymethylation be used as a marker for tumor diagnosis and treatment? Hum Genomics 14, 15 (2020). 10.1186/s40246-020-00265-5

34 Ramasamy, D. et al. Locus-Specific Enrichment Analysis of 5-Hydroxymethylcytosine Reveals Novel Genes Associated with Breast Carcinogenesis. Cells 11 (2022). 10.3390/cells11192939

35 Salmeron-Barcenas, E. G. et al. TET Enzymes and 5hmC Levels in Carcinogenesis and Progression of Breast Cancer: Potential Therapeutic Targets. Int J Mol Sci 25 (2023). 10.3390/ijms25010272

36 Zheng, Z., Ambigapathy, G. & Keifer, J. MeCP2 regulates Tet1-catalyzed demethylation, CTCF binding, and learning-dependent alternative splicing of the BDNF gene in Turtle. Elife 6 (2017). 10.7554/eLife.25384

37 Foong, Y. H. et al. TET1 displays catalytic and non-catalytic functions in the adult mouse cortex. Epigenetics 19, 2374979 (2024). 10.1080/15592294.2024.2374979

38 Marina, R. J. et al. TET-catalyzed oxidation of intragenic 5-methylcytosine regulates CTCF-dependent alternative splicing. EMBO J 35, 335–355 (2016). 10.15252/embj.201593235

39 Yadav, P. et al. Hypoxia-induced loss of SRSF2-dependent DNA methylation promotes CTCF-mediated alternative splicing of VEGFA in breast cancer. iScience 26, 106804 (2023). 10.1016/j.isci.2023.106804

40 Untergasser, A. et al. Primer3--new capabilities and interfaces. Nucleic Acids Res 40, e115 (2012). 10.1093/nar/gks596

41 Wang, S. & Liu, X. The UCSCXenaTools R package: a toolkit for accessing genomics data from UCSC Xena platform, from cancer multi-omics to single-cell RNA-seq. J Open Source Softw 4, 1627 (2019). 10.21105/joss.01627

42 Love, M. I., Huber, W. & Anders, S. Moderated estimation of fold change and dispersion for RNA-seq data with DESeq2. Genome Biol 15, 550 (2014). 10.1186/s13059-014-0550-8

43 Hulsen, T., de Vlieg, J. & Alkema, W. BioVenn - a web application for the comparison and visualization of biological lists using area-proportional Venn diagrams. BMC Genomics 9, 488 (2008). 10.1186/1471-2164-9-488

44 Wu, T. et al. clusterProfiler 4.0: A universal enrichment tool for interpreting omics data. Innovation (Camb) 2, 100141 (2021). 10.1016/j.xinn.2021.100141

