## Supplementary Data 1 for "Cytoplasmic localization of 5hmC reveals RNA hydroxymethylation in breast cancer"

**Supplementary Figure 1.** Hydroxymethylation in the total RNA and mRNA of BC cells, with technical positive control.
**Supplementary Figure 2.** 5mC is predominantly a nuclear feature of BC tissue.
**Supplementary Figure 3.** Levels of 5hmC are higher in colon cancer tissue.
**Supplementary Figure 4.** TET3 expression differences in the BCD cohort.
**Supplementary Figure 5.** Gene Set Enrichment Analysis associated with each TET in the BCD cohort.
**Supplementary Figure 6.** Spliceosome pathway visualization.
**Supplementary Figure 7.** Comparison of IHC quantification methods.
**Supplementary Figure 8.** TET expression in breast cancer and validation of TET3-regulated genes.
**Supplementary Figure 9.** Pearson correlation coefficients between TET1, TET2, TET3 and candidate target genes within breast cancer molecular subtypes.
**Supplementary Figure 10.** TET-family-associated nucleolar and small nucleolar/Cajal body RNA (snoRNA/scaRNA) gene signature in the BCD cohort.
**Supplementary Data 1**
**Supplementary Data 2**
**Supplementary Data 3**
**Supplementary Data 4**
**Supplementary Data 5**
**Supplementary Data 6.**
**Supplementary Data 7.**
**Supplementary Data 8.**

#### Suppl. Figure 1

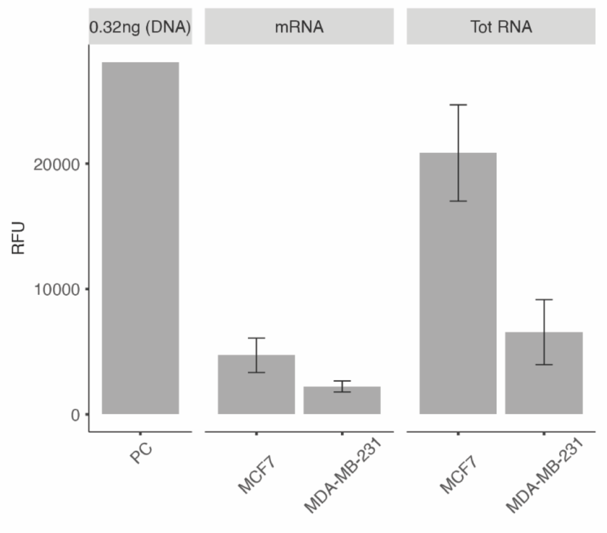


**Supplementary Figure 1. Hydroxymethylation in the total RNA and mRNA of BC cells, with technical positive control.** A commercially available control (PC = 0.32ng of 5hmC in the DNA) is used as a positive control to test the ability of the antibody to detect 5hmC. The bar chart shows the level of 5hmC content of total RNA and mRNA samples from MCF7 and MDA-MB-231 cells using an adaptation of the MethylFlash™ 5-mC RNA Methylation ELISA Easy Kit. Data are shown as relative fluorescence units (RFU). The error bars represent the standard deviation for n=3. The figure is descriptive and therefore does not include significance testing.

#### Suppl. Figure 2

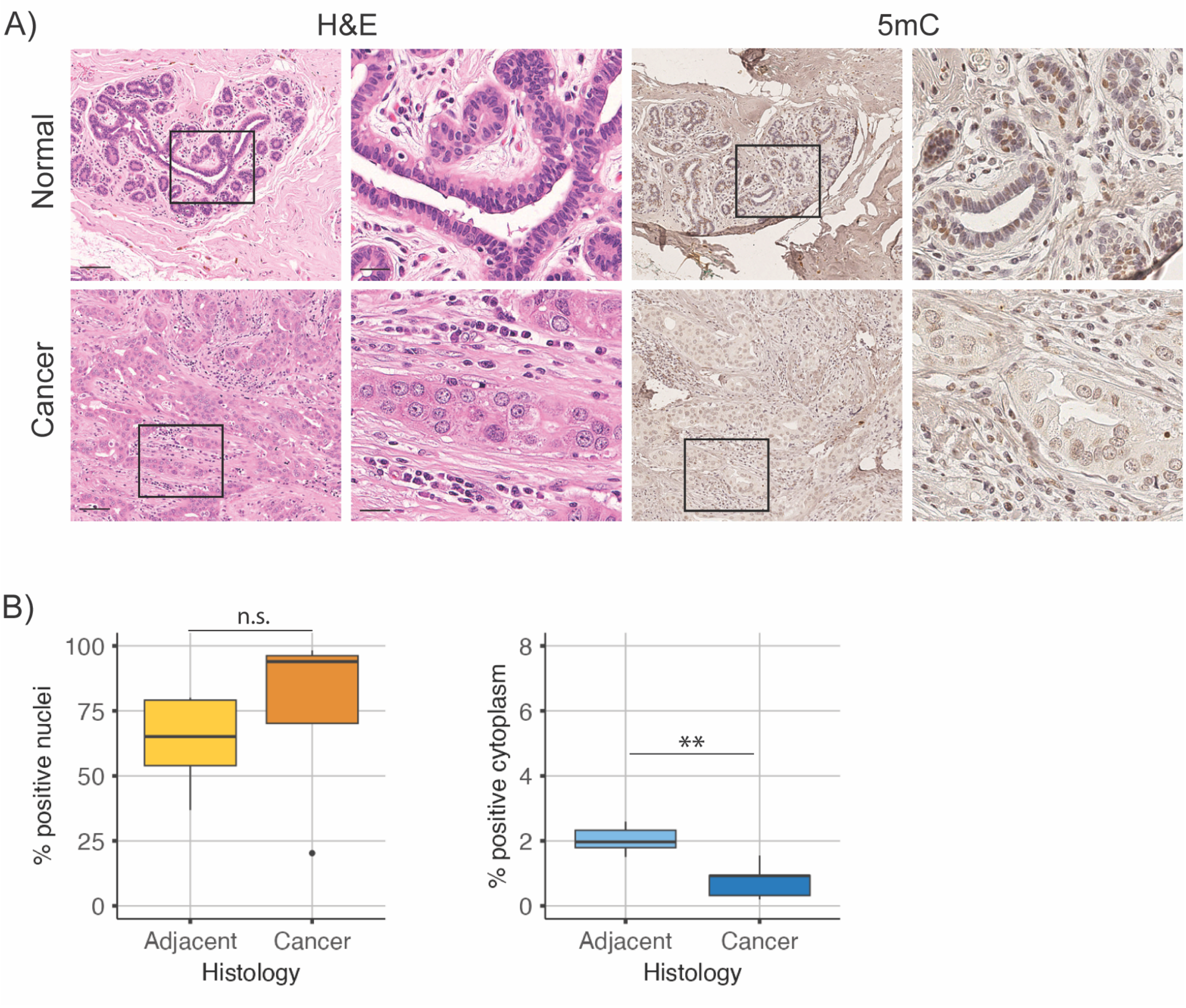


**Supplementary Figure 2. 5mC is predominantly a nuclear feature of BC tissue.** (A) H&E and 5mC immunohistochemistry of representative serial sections of non-affected adjacent breast tissue and the matching invasive ductal carcinoma. High-power images are shown in X) – Y). Scale bar=100µm; inset scale bar=25µm. (B) The nuclei of cancer cells contain significantly lower levels of 5mC compared to the non-affected counterpart, as estimated with a Welch Two Sample t-test. For both graphs, n=5 for each sample group. Boxplots display the interquartile range (IQR) and median. Significance is indicated as ** ($p<0.01$), or n.s. (non-significant).

#### Suppl. Figure 3

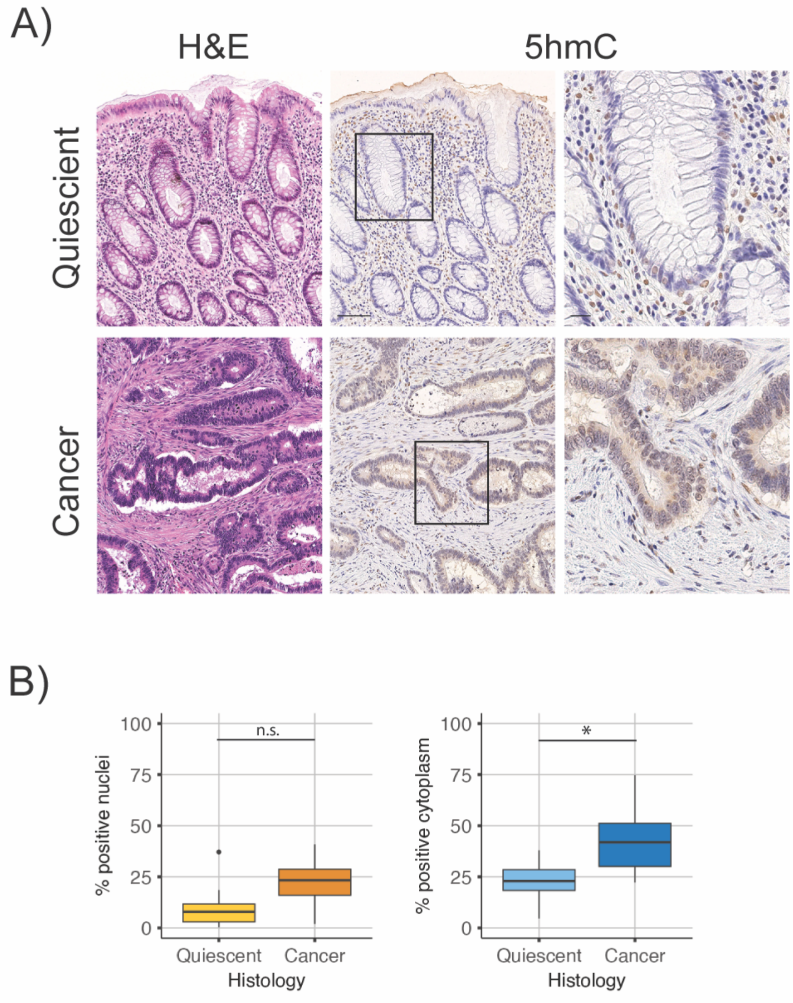


**Supplementary Figure 3. Levels of 5hmC are higher in colon cancer tissue.** (A) H&E and 5hmC immunohistochemistry of representative serial sections of a quiescent colon tissue from an IBD patient and the matching colitis-associated cancer. High-power images are shown in X) – Y). Scale bar=100µm; inset scale bar=20µm. (B) Both the nuclei and the cytoplasm of cancer cells contain a significantly higher levels of 5hmC compared to the normal counterpart, as estimated with a Welch Two Sample t-test. For both graphs, n=12 for each sample group. Boxplots display the interquartile range (IQR) and median. Significance is indicated as * (p<0.05), or n.s. (non-significant).

### Suppl. Figure 4

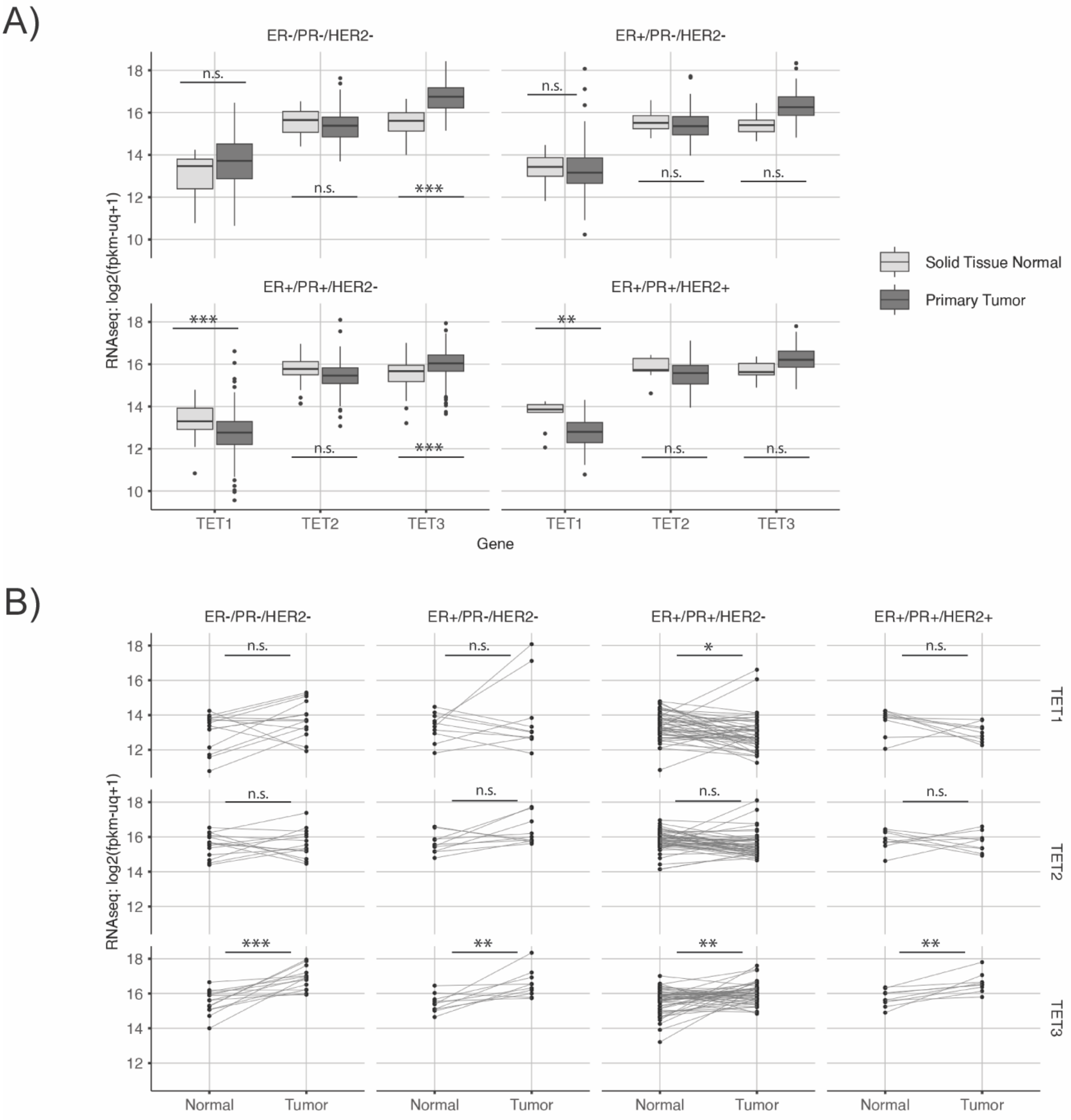


**Supplementary Figure 4**. **TET3 expression differences in subsets of BC samples.** (A) A significant increase in *TET3* mRNA expression is found in cancer samples with low *HER2* expression compared to the non-affected tissues, in both A) unpaired and B) paired analyses from the TCGA dataset. Boxplots display the interquartile range (IQR) and median. Significance is indicated as * ($p<0.05$), ** ($p < 0.01$), ***($p < 0.001$) or n.s. (non-significant).

#### Suppl. Figure 5

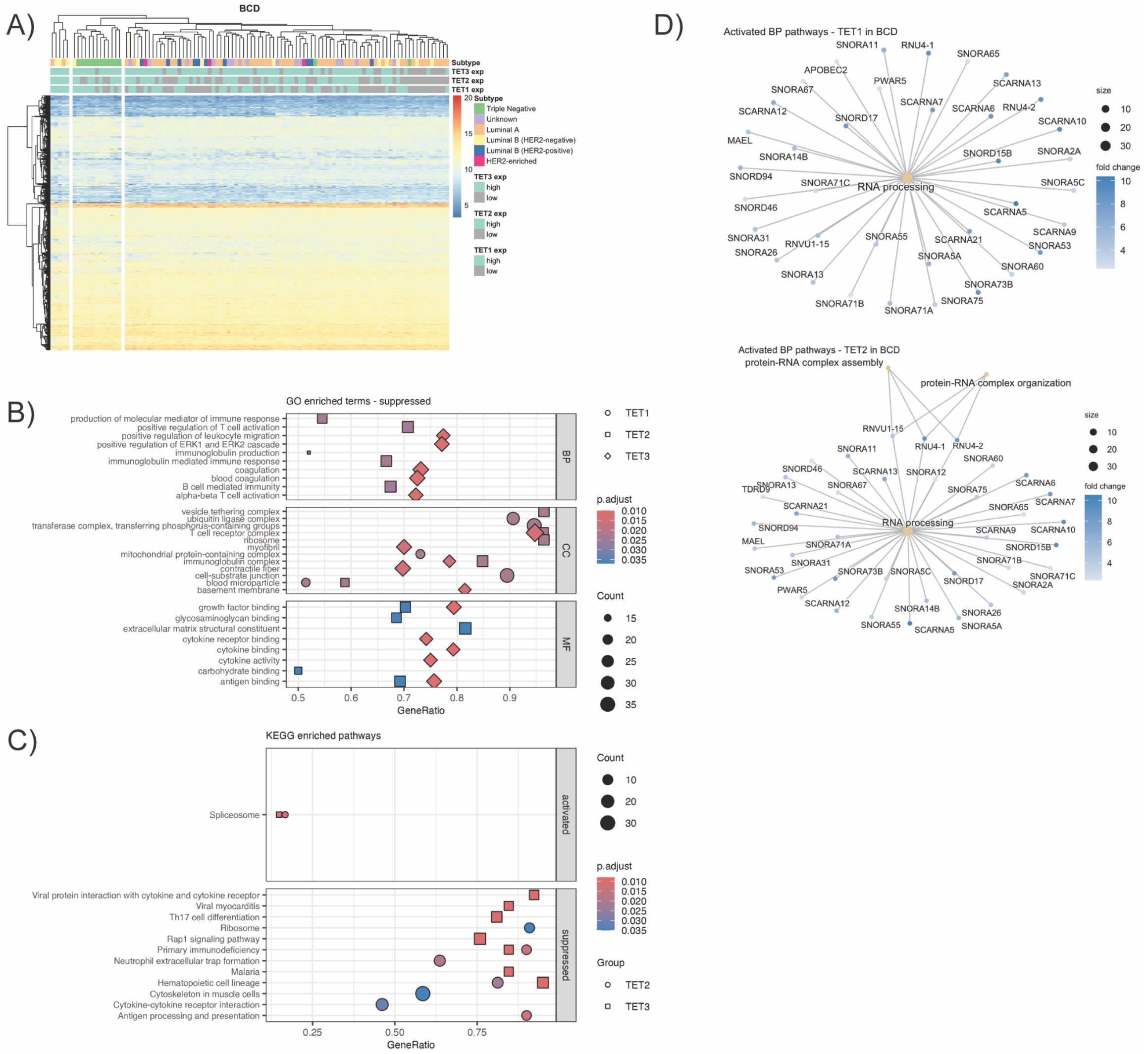


**Supplementary Figure 5. Gene Set Enrichment Analysis associated with each TET in the BCD cohort**. (A) Heatmap showing gene expression data for the BCD cohort, highlighting patterns across different subtypes and *TET* mRNA expression (TET_exp, defined as expression in cancer normalized to the matching normal). (B) gseGO enrichment results of DEGs (p adj < 0.05). A maximum of 5 significantly enriched and suppressed GO terms, including BP, CC, and MF, are shown for each TET group. (D) Category network plot displaying selected GO terms associated with TET1 and TET2 expression. The size of each category node represents the p-value of the GO term, with fold change values of the genes indicated by color intensity. (E) gseKEGG enrichment results of DEGs (p adj < 0.05). A maximum of 7 significantly enriched KEGG pathways are shown for each TET group.

#### Suppl. Figure 6

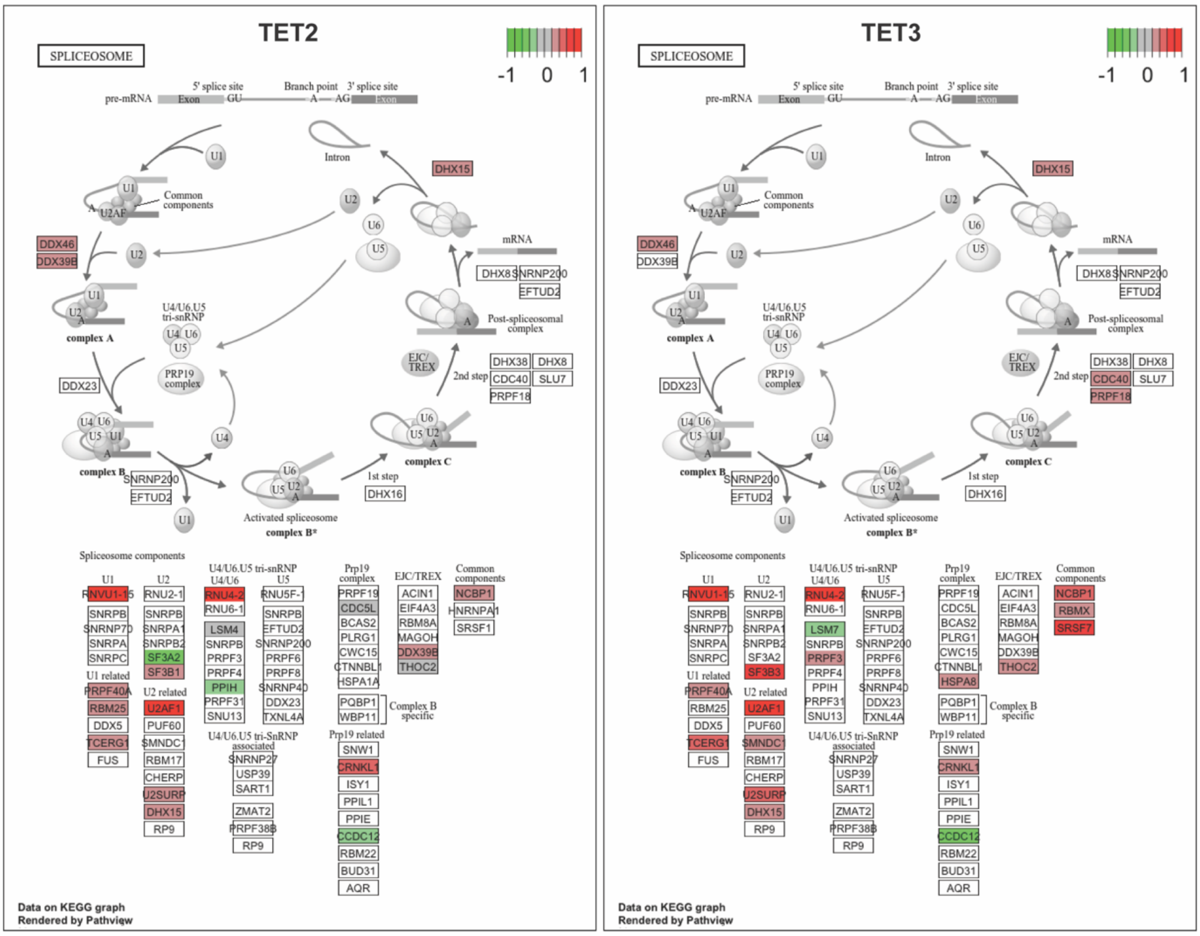


**Supplementary Figure 6. Spliceosome pathway visualization.** Comparative analysis of differential gene expression of the spliceosome pathway (hsa03040) in BC samples associated with TET2 (left) or TET3 (right) expression. Color highlights genes with differential expression following the scale of change log2FC.

**Suppl. Figure 7**

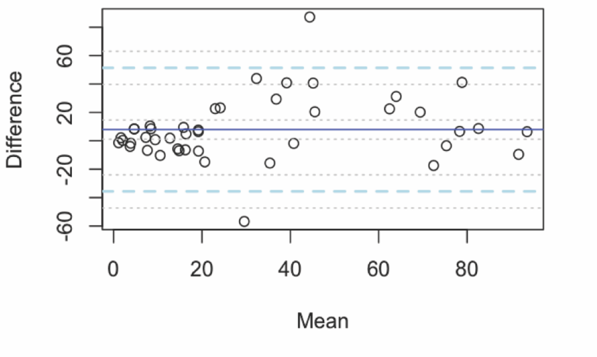


**Supplementary Figure 7. Comparison of IHC quantification methods.** The agreement between the two quantification methods employed (Visiopharm and QuPath) was assessed using a Bland and Altman plot on the cohort of breast tissues subjected to immunohistochemistry (IHC) for 5hmC. The limits of agreement, ranging from -1.96 standard deviations (sd) to +1.96 sd, are depicted with light blue lines, while the mean difference is represented by a solid blue line. Additionally, the 95% confidence interval is illustrated with grey lines.

#### Suppl. Figure 8

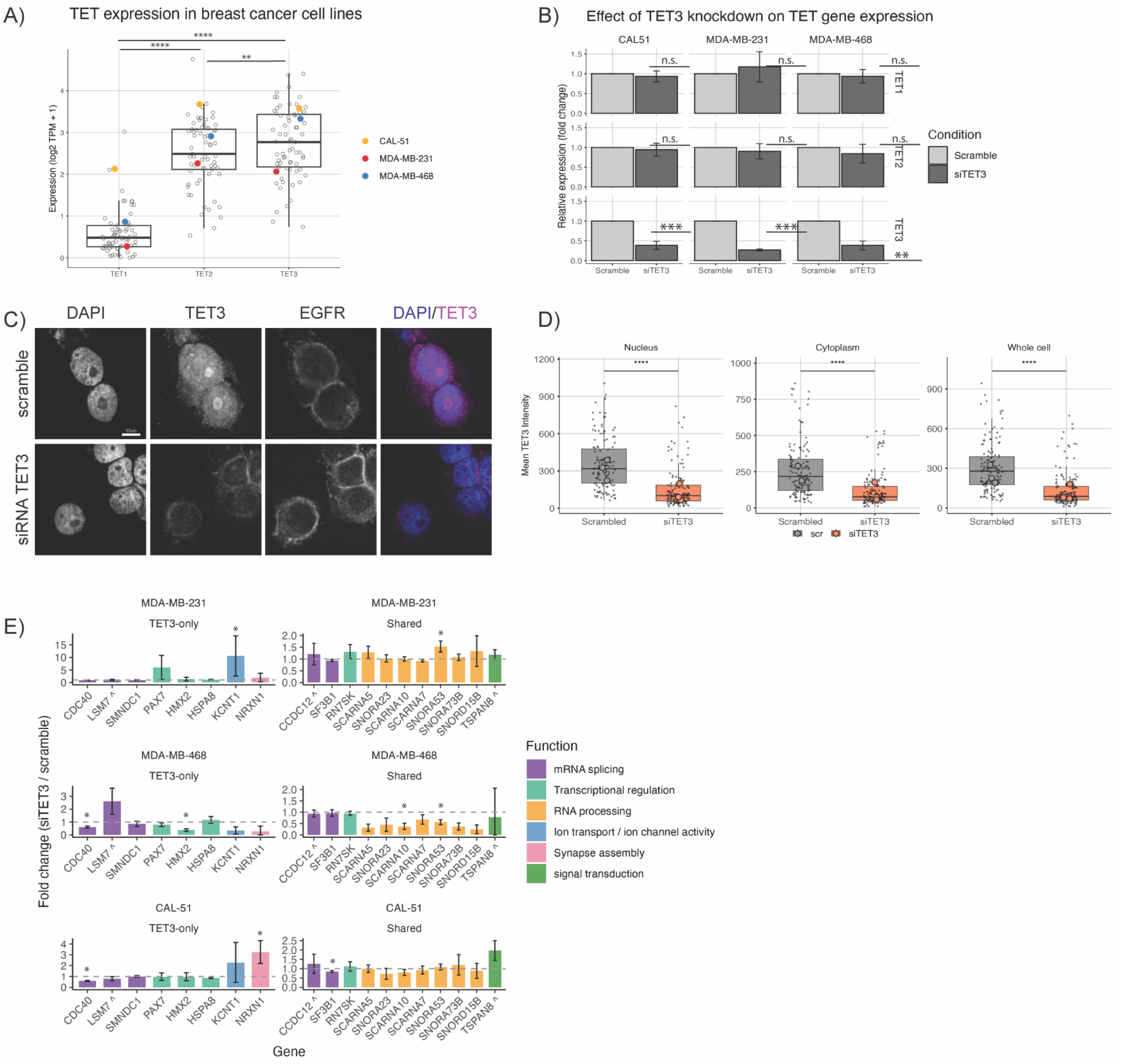


**Supplementary Figure 8. *TET* expression in breast cancer and validation of *TET3*-regulated genes.**

(A) Expression of *TET1*, *TET2*, and *TET3* across primary breast cancer cell lines from DepMap. Highlighted cell lines (MDA-MB-231, MDA-MB-468, CAL-51) were used for experimental validation. Data shown as $log_2$(TPM + 1). (B) Knockdown efficiency and specificity of siTET3 treatment in three TNBC cell lines. Expression measured by qPCR relative to scramble controls. Mean ± SD, n = 3-4 biological replicates. (C) Representative immunofluorescence images confirming TET3 knockdown at the protein level in MDA-MB-468 cells 48h after transfection with scrambled or TET3-targeting siRNA. DAPI (nucleus), TET3 and EGFR (used to delineate whole-cell boundaries) channels are shown individually and as a DAPI/TET3 merge. Scale bar = 10 µm. (D) Quantification of mean TET3 fluorescence intensity in the nucleus, cytoplasm and whole cell confirms a significant reduction in TET3 protein following siRNA knockdown relative to scrambled control. Linear mixed-effects model (log2(intensity) ~ Condition + (1|Replicate), fit on all cells, per compartment): nucleus, cytoplasm and whole cell, all $p < 0.0001$. Kruskal–Wallis test with Dunn's post-hoc on biological replicate medians (n = 3 per condition): nucleus $p = 0.050$ (*), whole cell $p = 0.050$ (*), cytoplasm $p = 0.127$ (n.s.). Boxplots display the IQR and median; small jittered points show individual cells (scrambled n = 136 cells; siTET3 n = 147 cells, pooled across 3 biological replicates); large outlined points represent biological replicate medians. (E) Fold-change in expression of selected candidate genes following *TET3* knockdown. Genes grouped by *TET3* specificity and colored by functional category. Genes marked with a caret (^) correspond to transcripts that exhibited a negative $\log_2$ fold change in the TCGA TET3-high versus TET3-low analysis. Dashed line indicates no change (fold-change = 1). Mean ± SD, n = 3-4 biological replicates. Statistical significance assessed by one-sample t-test on $\log_2$(fold-change) vs. 0. Significance is indicated as * ($p<0.05$), ** ($p < 0.01$), ***($p < 0.001$) or n.s. (non-significant). In Panel C), absence of stars indicates non-significancy.

#### Suppl. Figure 9

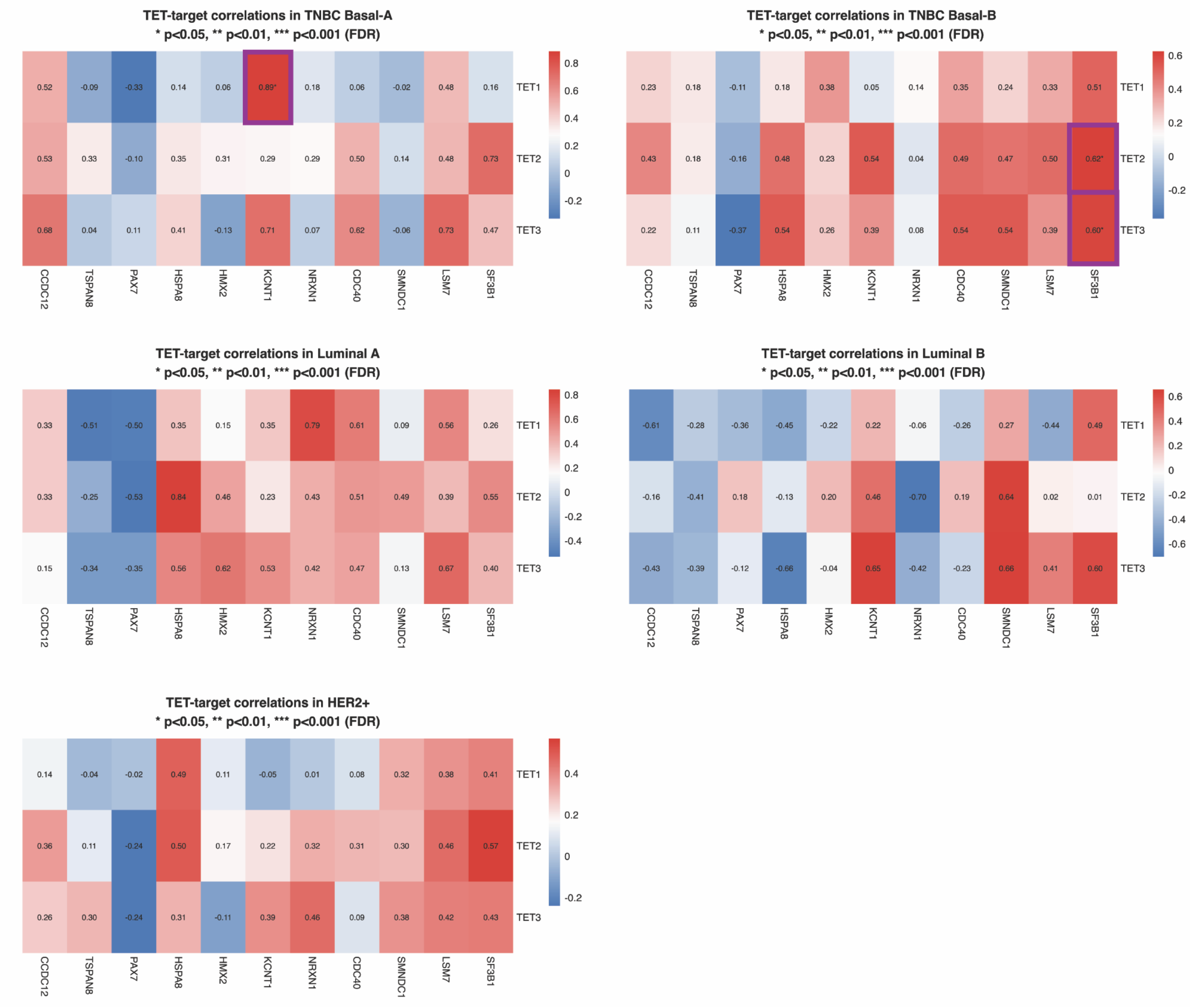


**Supplementary Figure 9. Pearson correlation coefficients between *TET1*, *TET2*, *TET3* and candidate target genes within breast cancer molecular subtypes.**
(First row) TNBC Basal-A (left, n = 9) and TNBC Basal-B (right, n = 19); (Second row) Luminal A (left, n = 10) and Luminal B (right, n = 8); (Third row) HER2+ (left, n = 17). Color intensity represents correlation strength (blue = negative, red = positive). Numbers show correlation coefficients with significance stars (* $p < 0.05$, ** $p < 0.01$, *** $p < 0.001$; FDR-adjusted, highlighted in purple). To maintain consistent multiple testing correction across visualization formats, FDR adjustment was performed on all pairwise gene correlations within each subtype. For clarity of presentation, heatmaps display only TET-target gene correlations (rectangular format), though significance testing accounts for the full correlation matrix. Data from Cancer Cell Line Encyclopedia (DepMap).

#### Suppl. Figure 10

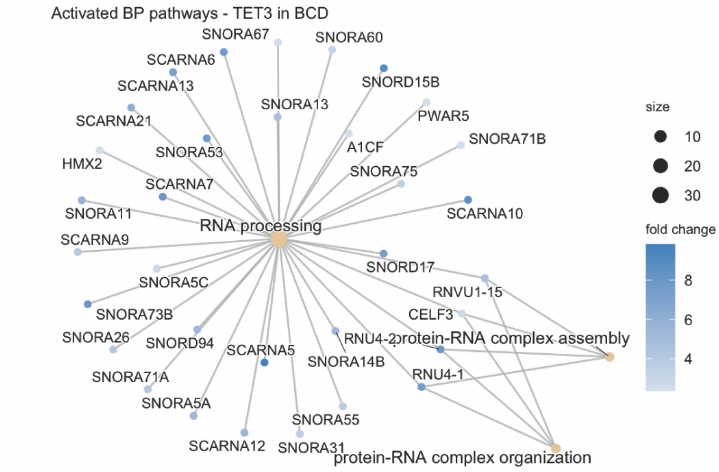


**Supplementary Figure 10. TET-family-associated nucleolar and small nucleolar/Cajal body RNA (snoRNA/scaRNA) gene signature in the BCD cohort.**
(A) Category network plot displaying selected GO terms associated with TET3 expression. The size of each category node represents the p-value of the GO term, and the fold change values of the genes are indicated by color intensity.

| Sample | Receptors | Grade |
|---|---|---|
| BC1 | ER+ PR+ HER2+ | 3 |
| BC3 | ER+ PR+ HER2+ | 2 |
| BC4 | ER+ PR+ HER2+ | 3 |
| BC5 | ER+ PR+ HER2+ | 3 |
| BC6 | ER+ PR+ HER2+ | 2 |
| BC7 | ER+ PR+ HER2+ | 2 |
| BC8 | ER+ PR+ HER2+ | 3 |
| BC9 | ER+ PR+ HER2- | 2 |
| BC10 | ER+ PR+ HER2- | 3 |
| BC11 | ER+ PR+ HER2- | 2 |
| BC13 | ER- PR- HER2+ | 3 |
| BC14 | ER- PR- HER2+ | 3 |
| BC16 | ER- PR- HER2+ | 3 |
| BC17 | ER- PR- HER2+ | 3 |
| BC18 | ER- PR- HER2- | 3 |
| BC19 | ER- PR- HER2- | 2 |
| BC20 | ER- PR- HER2- | 3 |
| BC21 | ER- PR- HER2- | 2 |
| BC22 | ER- PR- HER2- | 3 |
| BC23 | ER- PR- HER2- | NA |
| BC24 | ER- PR- HER2- | 2 |
| BC25 | ER- PR- HER2- | 3 |
| BC26 | ER- PR- HER2- | 3 |
| BC27 | ER- PR- HER2- | 3 |
| BC28 | ER- PR- HER2- | 3 |
| BC29 | ER+ PR- HER2- | 2 |

**Supplementary Data 1. Clinical information for BC samples analyzed in the study.**
Sample: Identifiers for each breast cancer sample.
Receptors: Expression status of estrogen receptor (ER), progesterone receptor (PR), and human epidermal growth factor receptor 2 (HER2) for each sample. "ER+" indicates positive expression, "ER-" indicates negative expression, and "NA" denotes missing data.
Stage: Clinical stage of breast cancer for each sample, including tumor (T), lymph nodes (N), and metastasis (M). The format follows the TNM staging system.
Grade: Histological grade of breast cancer for each sample, ranging from 1 to 3. "NA" denotes missing data.

##### a) Mean 5hmC Fluorescence Intensity (Nucleus vs Cytoplasm)

| Cell line | Median nuclear intensity | Median cytoplasmic intensity | LMM Estimate | SE | p value |
|---|---|---|---|---|---|
| BT474 | 54.500 | 35.300 | 0.678 | 0.074 | $4.87\times10^{-16}$ |
| MCF7 | 163.000 | 102.000 | 0.673 | 0.135 | $2.75\times10^{-6}$ |
| MDA-MB-231 | 46.300 | 42.100 | 0.325 | 0.074 | $2.15\times10^{-5}$ |
| SKBR3 | 105.000 | 74.800 | 0.360 | 0.054 | $2.61\times10^{-10}$ |

##### b) $Log_2$ Nuclear/Cytoplasmic Intensity Ratio

| Cell line | Median $log_2(N/C)$ | IQR $log_2(N/C)$ | LMM Estimate | SE | p value | Fold N/C |
|---|---|---|---|---|---|---|
| BT474 | 0.525 | 0.314 | 0.687 | 0.161 | 0.051 | 1.610 |
| MCF7 | 0.745 | 0.225 | 0.652 | 0.126 | 0.038 | 1.570 |
| MDA-MB-231 | 0.141 | 0.411 | 0.341 | 0.267 | 0.330 | 1.270 |
| SKBR3 | 0.340 | 0.107 | 0.348 | 0.054 | 0.020 | 1.270 |

Kruskal-Wallis across cell lines (on replicate medians): $\chi^2 = 4.385$, df = 3, p = 0.223 (n.s.)

| Dunn's pairwise post-hoc (Bonferroni-adjusted) | | |
|---|---|---|
| | Z | p adj |
| BT474 vs MCF7 | 0.226 | 1.000 |
| BT474 vs MDA-MB-231 | 1.580 | 0.678 |
| BT474 vs SKBR3 | 1.580 | 0.678 |
| MCF7 vs MDA-MB-231 | 1.360 | 1.000 |
| MCF7 vs SKBR3 | 1.360 | 1.000 |
| MDA-MB-231 vs SKBR3 | 0.000 | 1.000 |

##### c) Cytoplasmic Fraction of Total 5hmC Signal

| Cell line | Median F_cytoplasm | IQR F_cytoplasm | LMM Estimate | SE | p value | Fold N/C |
|---|---|---|---|---|---|---|
| BT474 | 0.443 (44.3%) | 0.008 | 0.438 | 0.022 | 0.118 | 1.270 |
| MCF7 | 0.477 (47.7%) | 0.049 | 0.496 | 0.024 | 0.890 | 1.270 |
| MDA-MB-231 | 0.504 (50.4%) | 0.118 | 0.443 | 0.065 | 0.478 | 1.610 |
| SKBR3 | 0.517 (51.7%) | 0.014 | 0.502 | 0.009 | 0.810 | 1.570 |

Kruskal-Wallis across cell lines (on replicate medians): $\chi^2 = 3.000$, df = 3, p = 0.392 (n.s.)

| Dunn's pairwise post-hoc (Bonferroni-adjusted) | | |
|---|---|---|
| | **Z** | **p adj** |
| BT474 vs MCF7 | −0.679 | 1.000 |
| BT474 vs MDA-MB-231 | −1.020 | 1.000 |
| BT474 vs SKBR3 | −1.700 | 0.537 |
| MCF7 vs MDA-MB-231 | −0.340 | 1.000 |
| MCF7 vs SKBR3 | −1.020 | 1.000 |
| MDA-MB-231 vs SKBR3 | −0.679 | 1.000 |

**Supplementary Data 2.** Statistical analyses of 5hmC fluorescence intensity in nuclear and cytoplasmic compartments of four BC cell lines (BT474, MCF7, MDA-MB-231, SKBR3), quantified from immunofluorescence images. (a) Raw mean fluorescence intensity by compartment. (b) $Log_2$ nuclear/cytoplasmic intensity ratio, testing whether nuclear 5hmC concentration exceeds cytoplasmic concentration. (c) Cytoplasmic fraction of total integrated 5hmC signal, testing whether the cytoplasm contributes more or less than 50% of total cellular signal when volume is accounted for. LMM: linear mixed-effects model with biological replicate as random intercept. ID: integrated density (mean intensity × area). In red, significant values are highlighted.

| a. Groups | Pr(>F) |
|---|---|
| TET1 | 0.023 |
| TET2 | <0.0001 |
| TET3 | <0.0001 |

| b. Tukey's multiple comparisons test | p adj | | |
|---|---|---|---|
| | TET1 | TET2 | TET3 |
| MCF7 vs BT474 | 0.026 | 0.036 | 0.001 |
| MDA-MB-231 vs BT474 | 0.086 | 0.000 | 0.000 |
| SKBR3 vs BT474 | 0.043 | 0.764 | 0.674 |
| MDA-MB-231 vs MCF7 | 0.822 | <0.0001 | <0.0001 |
| SKBR3 vs MCF7 | 0.980 | 0.143 | 0.005 |
| SKBR3 vs MDA-MB-231 | 0.960 | <0.0001 | <0.0001 |

**Supplementary Data 3**. a) Results of one-way analysis of variance (ANOVA) conducted to examine the differences in TET mRNA expression levels measured by qPCR among different cell lines. b) results of Tukey's post-hoc test, which compares the means of all pairs of groups to determine significant differences between them. Pr(>F): p-value ; p adj: adjusted p-value. In red, significant values are highlighted.

| Unpaired Welch two samples t-test | | | |
|---|---|---|---|
| 5hmC | | Sample size | |
| Nucleus | Cytoplasm | Adj | IDC |
| 0.535 | 0.003 | 23 | 24 |

| Paired t-test | | |
|---|---|---|
| 5hmC | | Sample size pairs |
| Nucleus | Cytoplasm | |
| 0.361 | 0.003 | 20 |

| Unpaired Welch two samples t-test | | | | |
|---|---|---|---|---|
| | 5hmC | | Sample size | |
| | Nucleus | Cytoplasm | Adj | IDC |
| ER- PR- HER2- | 0.808 | 0.012 | 10 | 10 |
| ER- PR- HER2+ | 0.169 | 0.124 | 4 | 4 |
| ER+ PR+ HER2+ | 0.706 | 0.483 | 7 | 7 |
| grade 2 | 0.881 | 0.257 | 6 | 6 |
| grade 3 | 0.275 | 0.01 | 14 | 15 |

| Paired t-test | | | |
|---|---|---|---|
| | 5hmC | | Sample size pairs |
| | Nucleus | Cytoplasm | |
| ER- PR- HER2- | 0.68 | 0.008 | 9 |
| ER- PR- HER2+ | 0.304 | 0.136 | 4 |
| ER+ PR+ HER2+ | 0.766 | 0.514 | 7 |
| grade 2 | 0.908 | 0.229 | 6 |
| grade 3 | 0.308 | 0.008 | 14 |

| Unpaired Welch two samples t-test | | | | |
|---|---|---|---|---|
| | | | Sample size | |
| | Nucleus | Cytoplasm | Adj | IDC |
| 5hmC | 0.391 | 0.004 | 5 | 6 |
| TET1 | 0.771 | 0.091 | | |
| TET2 | 0.355 | 0.018 | | |
| TET3 | 0.359 | 0.027 | | |

| Paired t-test | | | |
|---|---|---|---|
| | 5hmC | | Sample size pairs |
| | Nucleus | Cytoplasm | |
| 5hmC | 0.59 | 0.007 | 5 |
| TET1 | 0.989 | 0.206 | |
| TET2 | 0.442 | 0.023 | |
| TET3 | 0.263 | 0.027 | |

**Supplementary Data 4.** a-d) p value of paired and unpaired tests of the expression of 5hmC detected from IHC in normal-cancer pairs, including for samples divided by receptor status and grade. e-f) p value of paired and unpaired tests of the expression of 5hmC, TET1, TET2 and TET3 detected from IHC in normal-cancer pairs from ER-/PR- samples. In red, p value highlights the significant differences amongst groups.

| a. Two-way Anova | | | |
|---|---|---|---|
| | Pr(>F) | n normal | n tumor |
| All TETs | <0.0001 | 102 | 823 |

| b. Tukey's multiple comparisons test | | |
|---|---|---|
| | p adj | |
| | Primary tumor | Solid Tissue Normal |
| TET1 vs TET2 | <0.0001 | <0.0001 |
| TET1 vs TET3 | <0.0001 | <0.0001 |
| TET2 vs TET3 | <0.0001 | 0.858 |

| c. Tukey's multiple comparisons test | |
|---|---|
| Normal vs tumor | p adj |
| TET1 normal vs cancer | 0.007 |
| TET2 normal vs cancer | 0.046 |
| TET3 normal vs cancer | <0.0001 |

| d. Paired t-test | | | |
|---|---|---|---|
| p value | | | Sample size pairs |
| TET1 | TET2 | TET3 | |
| 0.382 | 0.786 | <0.0001 | 101 |

| e. One-way Anova | |
|---|---|
| | p value |
| TET1 | <0.0001 |
| TET2 | 0.056 |
| TET3 | <0.0001 |

| f. Tukey's multiple comparisons test | | | | | |
|---|---|---|---|---|---|
| | p adj | | | Sample size | |
| Normal vs tumor | TET1 | TET2 | TET3 | normal | tumor |
| ER- PR- HER2- | 0.110 | 0.992 | <0.0001 | 14 | 135 |
| ER+ PR- HER2- | >0.999 | 0.989 | 0.053 | 10 | 59 |
| ER+ PR+ HER2- | <0.0001 | 0.063 | 0.0003 | 50 | 340 |
| ER+ PR+ HER2+ | 0.004 | 0.792 | 0.214 | 9 | 71 |

| g. Paired t-test | | | | |
|---|---|---|---|---|
| | p value | | | Sample size pairs |
| Normal vs tumor | TET1 | TET2 | TET3 | |
| ER- PR- HER2- | 0.078 | 0.797 | 0.0004 | 14 |
| ER+ PR- HER2- | 0.476 | 0.069 | 0.006 | 10 |
| ER+ PR+ HER2- | 0.015 | 0.091 | 0.005 | 49 |
| ER+ PR+ HER2+ | 0.119 | 0.754 | 0.003 | 9 |

**Supplementary Data 5.** Statistical analyses of TET mRNA expression patterns in BC samples derived from The Cancer Genome Atlas Program (TCGA). a) the comparative analyses between cancerous and adjacent tissue samples, featuring statistical results from two-way ANOVA, b-c) Tukey's multiple comparisons test, and d) paired t-tests for primary tumor and solid tissue normal samples. e-g) subtype-specific analyses of TET mRNA expression, stratified by receptor status, and include statistical results from e) one-way ANOVA, f) Tukey's multiple comparisons test, and g) paired t-tests, offering insights into the differential expression patterns of TET mRNAs across various BC subtypes. Pr(>F): p-value; p adj: adjusted p-value; Adj: Adjacent non-affected tissue; IDC: invasive ductal carcinoma; ER: Estrogen receptor; PR: Progesterone receptor; HER2: human epidermal growth factor receptor 2. Values <0.05 are highlighted in red

| Target | Primer sequence (5’-3’) |
| --- | --- |
| TET1 | Fwd- AACGGAATCCAACGGGAAGG<br>Rev- TGCCATCACGTTAGCACACT |
| TET2 | Fwd- GATAGAACCAACCATGTTGAGGG<br>Rev-TGGAGCTTTGTAGCCAGAGGT |
| TET3 | Fwd-TCCAGCAACTCCTAGAACTGAG<br>Rev-AGGCCGCTTGAATACTGACTG |
| HPRT1 | Fwd- TGACACTGGCAAAACAATGCA<br>Rev-GGTCCTTTTCACCAGCAAGCT |

| Target | siRNA sequence (5’-3’) |
| --- | --- |
| TET3 | AAACAAAGCAAGUCGAGAA |
| | GGGCAAGAUUCCCGAGGAU |
| | CAUAUUAUAGCAACGGAAA |
| | ACGAGGAAAUAGCGAUUGA |
| scramble | UGGUUUACAUGUCGACUAA |
| | UGGUUUACAUGUUGUGUGA |
| | UGGUUUACAUGUUUUCUGA |
| | UGGUUUACAUGUUUUCCUA |

**Supplementary Data 6**. qPCR and siRNA sequences used in this study (5′–3′).

**a) Linear mixed-effects model on all cells**

| | LMM Estimate | SE | p-value | Fold-change (siTET3/scr) |
|---|---|---|---|---|
| Nucleus | −1.43 | 0.107 | $7.79 \times 10^{-32}$ | 0.370 |
| Cytoplasm | −1.18 | 0.119 | $3.04 \times 10^{-20}$ | 0.440 |
| Whole cell | −1.30 | 0.108 | $2.60 \times 10^{-27}$ | 0.410 |

**b) Kruskal–Wallis test with Dunn's post-hoc (Bonferroni-adjusted)**

| | $\chi^2$ | df | KW p-value | Dunn's Z | Dunn's p adj |
|---|---|---|---|---|---|
| Nucleus | 3.857 | 1.000 | 0.050 | 1.960 | 0.050 |
| Cytoplasm | 2.333 | 1.000 | 0.127 | 1.530 | 0.127 |
| Whole cell | 3.857 | 1.000 | 0.050 | 1.960 | 0.050 |

**Supplementary Data 7.** Statistical analyses of TET3 fluorescence intensity in nuclear, cytoplasmic and whole-cell compartments of MDA-MB-468 cells 48 h after transfection with scrambled (scr) or TET3-targeting (siTET3) siRNA, quantified from immunofluorescence images. (a) Linear mixed-effects model (log2(intensity) ~ Condition + (1|Replicate)) fit on all cells within each compartment, testing whether TET3 intensity differs between scrambled and siTET3 conditions. (b) Kruskal-Wallis test with Dunn's post-hoc (Bonferroni-adjusted) on biological replicate medians (n = 3 per condition), used as a rank-based cross-check of (a). LMM: linear mixed-effects model. In red, significant values are highlighted.

| Target Gene | TET Gene | Significant | N | Result |
|---|---|---|---|---|
| CCDC12 | TET1 | ns | 70 | r = 0.148 [-0.090, 0.370], p = 2.21e-01 |
| CCDC12 | TET2 | * | 70 | r = 0.289 [0.058, 0.491], p = 1.51e-02 |
| CCDC12 | TET3 | ns | 70 | r = 0.233 [-0.002, 0.444], p = 5.22e-02 |
| TSPAN8 | TET1 | ns | 70 | r = -0.012 [-0.247, 0.223], p = 9.19e-01 |
| TSPAN8 | TET2 | ns | 70 | r = 0.144 [-0.094, 0.367], p = 2.34e-01 |
| TSPAN8 | TET3 | ns | 70 | r = 0.084 [-0.154, 0.313], p = 4.88e-01 |
| PAX7 | TET1 | ns | 70 | r = -0.194 [-0.410, 0.043], p = 1.08e-01 |
| PAX7 | TET2 | * | 70 | r = -0.251 [-0.459, -0.017], p = 3.60e-02 |
| PAX7 | TET3 | ns | 70 | r = -0.232 [-0.443, 0.003], p = 5.36e-02 |
| HSPA8 | TET1 | ns | 70 | r = 0.197 [-0.039, 0.413], p = 1.01e-01 |
| HSPA8 | TET2 | *** | 70 | r = 0.550 [0.362, 0.695], p = 7.96e-07 |
| HSPA8 | TET3 | *** | 70 | r = 0.463 [0.255, 0.629], p = 5.55e-05 |
| HMX2 | TET1 | ns | 70 | r = 0.038 [-0.198, 0.271], p = 7.52e-01 |
| HMX2 | TET2 | ns | 70 | r = -0.011 [-0.246, 0.224], p = 9.25e-01 |
| HMX2 | TET3 | ns | 70 | r = 0.069 [-0.169, 0.299], p = 5.73e-01 |
| KCNT1 | TET1 | ns | 70 | r = 0.158 [-0.080, 0.379], p = 1.90e-01 |
| KCNT1 | TET2 | ** | 70 | r = 0.381 [0.160, 0.565], p = 1.15e-03 |
| KCNT1 | TET3 | ** | 70 | r = 0.317 [0.089, 0.514], p = 7.50e-03 |
| NRXN1 | TET1 | ns | 70 | r = 0.154 [-0.084, 0.376], p = 2.02e-01 |
| NRXN1 | TET2 | ns | 70 | r = -0.010 [-0.245, 0.225], p = 9.32e-01 |
| NRXN1 | TET3 | ns | 70 | r = 0.068 [-0.170, 0.298], p = 5.77e-01 |
| CDC40 | TET1 | * | 70 | r = 0.251 [0.017, 0.459], p = 3.61e-02 |
| CDC40 | TET2 | *** | 70 | r = 0.505 [0.306, 0.661], p = 8.28e-06 |
| CDC40 | TET3 | *** | 70 | r = 0.447 [0.237, 0.617], p = 1.04e-04 |
| SMNDC1 | TET1 | * | 70 | r = 0.256 [0.023, 0.463], p = 3.22e-02 |
| SMNDC1 | TET2 | *** | 70 | r = 0.475 [0.270, 0.639], p = 3.24e-05 |
| SMNDC1 | TET3 | *** | 70 | r = 0.404 [0.187, 0.584], p = 5.15e-04 |
| LSM7 | TET1 | * | 70 | r = 0.303 [0.073, 0.502], p = 1.08e-02 |
| LSM7 | TET2 | ** | 70 | r = 0.344 [0.119, 0.536], p = 3.50e-03 |
| LSM7 | TET3 | *** | 70 | r = 0.401 [0.183, 0.581], p = 5.84e-04 |
| SF3B1 | TET1 | *** | 70 | r = 0.399 [0.181, 0.579], p = 6.31e-04 |
| SF3B1 | TET2 | *** | 70 | r = 0.535 [0.343, 0.684], p = 1.83e-06 |
| SF3B1 | TET3 | *** | 70 | r = 0.496 [0.295, 0.654], p = 1.29e-05 |
| TET1 | TET1 | *** | 70 | r = 1.000 [1.000, 1.000], p = 0.00e+00 |
| TET1 | TET2 | *** | 70 | r = 0.480 [0.276, 0.642], p = 2.62e-05 |
| TET1 | TET3 | *** | 70 | r = 0.482 [0.279, 0.644], p = 2.37e-05 |
| TET2 | TET1 | *** | 70 | r = 0.480 [0.276, 0.642], p = 2.62e-05 |
| TET2 | TET2 | *** | 70 | r = 1.000 [1.000, 1.000], p = 0.00e+00 |
| TET2 | TET3 | *** | 70 | r = 0.736 [0.606, 0.828], p = 3.98e-13 |
| TET3 | TET1 | *** | 70 | r = 0.482 [0.279, 0.644], p = 2.37e-05 |
| TET3 | TET2 | *** | 70 | r = 0.736 [0.606, 0.828], p = 3.98e-13 |
| TET3 | TET3 | *** | 70 | r = 1.000 [1.000, 1.000], p = 0.00e+00 |

**Supplementary Data 8.** Correlation of selected genes with TET family expression in DepMap Shown are Pearson correlation coefficients (r), 95% confidence intervals (in square brackets), two-sided Benjamini–Hochberg adjusted p-values, significance codes (Significant: ns, * < 0.05, ** < 0.01, *** < 0.001), and sample size (N).
