## Supplementary Data 2 for "Cytoplasmic localization of 5hmC reveals RNA hydroxymethylation in breast cancer"

**Differentially Expressed Genes uniquely in TET3-high vs TET-low samples**

| Gene | Primer | Primer sequence (5’-3’) | Condition | $log_2FC$ | adj p-value | Involved biological processes (BP, QuickGO) | Selection Category (GSEA via ClusterProfiler) |
|---|---|---|---|---|---|---|---|
| **PAX7** | Fwd<br>Rev | ACCCCTGCCTAACCACATC<br>GCGGCAAAGAATCTTGGAGAC | TET3-high vs low | 6.937 | 1.50E-08 | GO:0006355 (regulation of DNA-templated transcription), **GO:0006357 (regulation of transcription by RNA polymerase II)**, GO:0007399 (nervous system development), GO:0007517 (muscle organ development), GO:0009653 (anatomical structure morphogenesis), GO:0014816 (skeletal muscle satellite cell differentiation), GO:0043066 (negative regulation of apoptotic process) | Top-ranked by effect size |
| **HMX2** | Fwd<br>Rev | GCTTCTGCCCAGACCAGCACG<br>AGTGCGAAGGAGAGAGGAAAGG | TET3-high vs low | 2.345 | 2.35E-02 | **GO:0006355 (regulation of DNA-templated transcription), GO:0006357 (regulation of transcription by RNA polymerase II)**, GO:0007399 (nervous system development), GO:0030154 (cell differentiation) | GO-derived (RNA processing) |
| **KCNT1** | Fwd<br>Rev | CATTGTCTATCTCATCCGCTCC<br>TGAGTAGAGTGTGCCATCCC | TET3-high vs low | 2.150 | 8.06E-03 | GO:0006811 (monoatomic ion transport), GO:0006813 (potassium ion transport), GO:0034220 (monoatomic ion transmembrane transport), GO:0051289 (protein homotetramerization), GO:0071805 (potassium ion transmembrane transport) | GO-derived (potassium ion transmembrane transport) |
| **NRXN1** | Fwd<br>Rev | TAAGTGGCCTCCTAATGACCG<br>TCGCACCAATACGGCTTCTTT | TET3-high vs low | 2.111 | 7.01E-03 | GO:0001525 (angiogenesis), GO:0007155 (cell adhesion), GO:0007157 (heterophilic cell-cell adhesion), GO:0007158 (neuron cell-cell adhesion), GO:0007165 (signal transduction), GO:0007399 (nervous system development), GO:0007416 (synapse assembly), GO:0007612 (learning), GO:0010628 (positive regulation of gene expression), GO:0010629 (negative regulation of gene expression), GO:0016339 (calcium-dependent cell-cell adhesion), GO:0021707 (cerebellar granule cell differentiation), GO:0023041 (neuronal signal transduction), GO:0030534 (adult behavior), GO:0031175 (neuron projection development), GO:0035176 (social behavior), GO:0035418 (protein localization to synapse), GO:0045184 (establishment of protein localization), GO:0045743 (positive regulation of fibroblast growth factor receptor signaling pathway), GO:0051490 (negative regulation of filopodium assembly), GO:0051897 (positive regulation of phosphatidylinositol 3-kinase/protein kinase B signal transduction), GO:0051965 (positive regulation of synapse assembly), GO:0051968 (positive regulation of synaptic transmission, glutamatergic), GO:0070374 (positive regulation of ERK1 and ERK2 cascade), GO:0071277 (cellular response to calcium ion), GO:0071625 (vocalization behavior), GO:0090126 (protein-containing complex assembly involved in synapse maturation), GO:0097091 (synaptic vesicle clustering), GO:0097104 (postsynaptic membrane assembly), GO:0097105 (presynaptic membrane assembly), GO:0097112 (gamma-aminobutyric acid receptor clustering), GO:0097114 (NMDA glutamate receptor clustering), GO:0097116 (gephyrin clustering involved in postsynaptic density assembly), GO:0097117 (guanylate kinase-associated protein clustering), GO:0097118 (neuroligin clustering involved in postsynaptic membrane assembly), GO:0097119 (postsynaptic density protein 95 clustering), GO:0097120 (receptor localization to synapse), GO:0106071 (positive regulation of adenylate cyclase-activating G protein-coupled receptor signaling pathway), GO:0141163 (positive regulation of cAMP/PKA signal transduction), GO:1900738 (positive regulation of phospholipase C-activating G protein-coupled receptor signaling pathway), GO:1903078 (positive regulation of protein localization to plasma membrane), GO:1905520 (positive regulation of presynaptic active zone assembly), GO:2000463 (positive regulation of excitatory postsynaptic potential) | GO-derived (regulation of membrane potential, neurotransmitter transport) |
| **HSPA8** | Fwd<br>Rev | TCCTACCAAGCAGACACAGACC<br>CAGGAGGTATGCCTGTGAGTTC | TET3-high vs low | 0.356 | 4.85E-02 | **GO:0000398 (mRNA splicing, via spliceosome), GO:0006397 (mRNA processing)**, GO:0006457 (protein folding), GO:0006914 (autophagy), GO:0006986 (response to unfolded protein), GO:0007165 (signal transduction), **GO:0008380 (RNA splicing)**, GO:0009267 (cellular response to starvation), GO:0009968 (negative regulation of signal transduction), GO:0030335 (positive regulation of cell migration), GO:0031647 (regulation of protein stability), GO:0033554 (cellular response to stress), GO:0039531 (regulation of cytoplasmic pattern recognition receptor signaling pathway), GO:0042026 (protein refolding), **GO:0043254 (regulation of protein-containing complex assembly), GO:0045892 (negative regulation of DNA-templated transcription)**, GO:0046034 (ATP metabolic process), GO:0051129 (negative regulation of cellular component organization), GO:0061024 (membrane organization), **GO:0061635 (regulation of protein complex stability)**, GO:0061684 (chaperone-mediated autophagy), GO:0061740 (protein targeting to lysosome involved in chaperone-mediated autophagy), GO:0071383 (cellular response to steroid hormone stimulus), GO:0071383 (cellular response to steroid hormone stimulus), GO:0072318 (clathrin coat disassembly), GO:0160020 (positive regulation of ferroptosis), GO:1900226 (negative regulation of NLRP3 inflammasome complex assembly), GO:1902903 (regulation of supramolecular fiber organization), GO:1902904 (negative regulation of supramolecular fiber organization), GO:1904589 (regulation of protein import), GO:1904764 (chaperone-mediated autophagy translocation complex disassembly) | KEGG (spliceosome) |
| **CDC40** | Fwd<br>Rev | GAGACTGGAGTTCACCTTGACC<br>GGCAGCCATTTGCTGTGTCCTA | TET3-high vs low | 0.330 | 2.82E-02 | **GO:0000398 (mRNA splicing, via spliceosome), GO:0006397 (mRNA processing), GO:0008380 (RNA splicing),** GO:1990403 (embryonic brain development) | KEGG (spliceosome) |
| **SMNDC1** | Fwd<br>Rev | GGCAGTCTGGAGTGAAGATGGA<br>GGCTTGAGGTTCAACAGTGGAG | TET3-high vs low | 0.240 | 1.80E-02 | **GO:0000375 (RNA splicing, via transesterification reactions), GO:0006397 (mRNA processing)**, GO:0006915 (apoptotic process), **GO:0008380 (RNA splicing)** | KEGG (spliceosome) |
| **LSM7** | Fwd<br>Rev | CACTCCTCAACCTTGTGCTGGA<br>GCAGATTAGCACCACGGACGTG | TET3-high vs low | -0.335 | 4.76E-02 | **GO:0000244 (spliceosomal tri-snRNP complex assembly), GO:0000290 (deadenylation-dependent decapping of nuclear-transcribed mRNA), GO:0000387 (spliceosomal snRNP assembly), GO:0000398 (mRNA splicing, via spliceosome), GO:0000956 (nuclear-transcribed mRNA catabolic process), GO:0006397 (mRNA processing), GO:0008380 (RNA splicing)** | KEGG (spliceosome) |

**Differentially Expressed Genes associated with multiple TETs**

| Gene | Primer | Primer sequence (5’-3’) | Condition | $log_2FC$ | adj p-value | Involved biological processes (BP, QuickGO) or reported function (NCBI Gene) | Selection Category (GSEA via ClusterProfiler) |
|---|---|---|---|---|---|---|---|
| **CCDC12** | Fwd | CCTGACTGGGACCTCAAGAG | TET3-high vs low | -0.436 | 7.17E-03 | Part of the U2-type spliceosomal complex and post-mRNA release spliceosomal complex, functioning in **pre-mRNA splicing.** | KEGG (spliceosome) |
| | Rev | CAGTCGGAGTCACAGGTCTT | TET2-high vs low | -0.309 | 2.18E-02 | | |
| **SF3B1** | Fwd | TTTGCTGGATACGTGACATCAA | TET3-high vs low | 0.241 | 5.46E-03 | **GO:0000398 (mRNA splicing, via spliceosome), GO:0000245 (spliceosomal complex assembly), GO:0000375 (RNA splicing, via transesterification reactions), GO:0000387 (spliceosomal snRNP assembly), GO:0000388 (spliceosome conformational change to release U4 (or U4atac) and U1 (or U11))**, GO:0006338 (chromatin remodeling), **GO:0006397 (mRNA processing), GO:0008380 (RNA splicing), GO:0045943 (positive regulation of transcription by RNA polymerase I), GO:0045944 (positive regulation of transcription by RNA polymerase II), GO:0045945 (positive regulation of transcription by RNA polymerase III), GO:1903241 (U2-type prespliceosome assembly)** | KEGG (spliceosome) |
| | Rev | CCGGTCTGCAATCTTTGGAGG | TET2-high vs low | 0.246 | 1.97E-04 | | |
| **RN7SK** | Fwd | CAAACAAGCTCTCAAGGTCCA | TET3-high vs low | 8.915 | 6.32E-14 | Predicted to be involved in **negative regulation of transcription elongation by RNA polymerase II**. Part of 7SK snRNP. | Top-ranked by effect size |
| | Rev | GCCTCATTTGGATGTGTCTG | TET1-high vs low | 9.535 | 4.60E-39 | | |
| | | | TET2-high vs low | 9.604 | 9.12E-41 | | |
| **SCARNA10** | Fwd | GTTTGGCTAAGCCCAGGGAC | TET3-high vs low | 8.647 | 6.33E-13 | Predicted to be involved in **RNA processing**. Located in Cajal body. | Top-ranked by effect size; GO-derived (RNA processing) |
| | Rev | GTTGGTCTGCCCTTACAGTGA | TET1-high vs low | 9.197 | 1.65E-31 | | |
| | | | TET2-high vs low | 9.096 | 4.09E-28 | | |
| **SCARNA5** | Fwd | TGAATGTCACGGTCCCTTTGTTGC | TET3-high vs low | 9.816 | 2.74E-12 | Predicted to be involved in **RNA processing**. Located in Cajal body. | Top-ranked by effect size |
| | Rev | AGCTGCTCCATGATCCCATACACA | TET1-high vs low | 10.354 | 6.25E-32 | | |
| | | | TET2-high vs low | 10.518 | 5.97E-33 | | |
| **SNORA73B** | Fwd | AGGCTCTGTCCAAGTGGCATAGG | TET3-high vs low | 8.004 | 2.74E-12 | Predicted to be involved in **RNA processing**. Predicted to be located in nucleolus. | Top-ranked by effect size; GO-derived (RNA processing) |
| | Rev | CGAGGCCCAGCTTCATCTTCAAC | TET1-high vs low | 8.726 | 1.10E-32 | | |
| | | | TET2-high vs low | 8.624 | 3.20E-29 | | |
| **SCARNA7** | Fwd | ACTGACTGGTGATGCTGATATT | TET3-high vs low | 8.211 | 4.88E-11 | Predicted to be involved in **RNA processing**. Predicted to be located in nucleolus. | Top-ranked by effect size; GO-derived (RNA processing) |
| | Rev | GCTAGTTAGGTGAGCTGTTCTT | TET1-high vs low | 7.985 | 1.10E-32 | | |
| | | | TET2-high vs low | 8.619 | 1.15E-26 | | |
| **SNORA53** | Fwd | ACATGCTTCCTTAGATCCACCT | TET3-high vs low | 7.714 | 4.88E-11 | Predicted to be involved in **RNA processing**. Predicted to be located in nucleolus. | Top-ranked by effect size; GO-derived (RNA processing) |
| | Rev | CACACGAGTCGAGTCCTGAT | TET1-high vs low | 8.147 | 3.36E-30 | | |
| | | | TET2-high vs low | 8.209 | 1.25E-28 | | |
| **TSPAN8** | Fwd | CATAGGCTTGCTTCTGATCCTGC | TET3-high vs low | -0.416 | 1.49E-11 | GO:0030195 (negative regulation of blood coagulation) | Top-ranked by effect size |
| | Rev | CTTTCCCCTGTGGCGCTCAAAA | TET1-high vs low | 0.195 | 1.41E-05 | | |
| | | | TET2-high vs low | 0.052 | 3.13E-04 | | |
| **SNORA23** | Fwd | TCATGCGGCCAAAGAGTAAC | TET3-high vs low | 8.851 | 6.43E-10 | Predicted to be involved in **RNA processing**. Predicted to be located in nucleolus. | Top-ranked by effect size |
| | Rev | GGCCAGTGGTAGATGTGTCC | TET1-high vs low | 9.020 | 1.03E-21 | | |
| | | | TET2-high vs low | 8.977 | 8.72E-20 | | |
| **SNORD15B** | Fwd | ACGAGTCAGAAAGGTCACGT | TET3-high vs low | 8.937 | 4.49E-08 | Predicted to be involved in **RNA processing**. Predicted to be located in nucleolus. | Top-ranked by effect size; GO-derived (RNA processing) |
| | Rev | CCTTCTCAGACAAATGCCTCT | TET1-high vs low | 9.477 | 4.57E-20 | | |
| | | | TET2-high vs low | 9.389 | 6.38E-18 | | |
